# Structural insights into the peptide selectivity and activation of human neuromedin U receptors

**DOI:** 10.1101/2022.02.21.481304

**Authors:** Chongzhao You, Yumu Zhang, Peiyu Xu, Sijie Huang, Wanchao Yin, H. Eric Xu, Yi Jiang

## Abstract

Neuromedin U receptors (NMURs), including NMUR1 and NMUR2, are a group of G_q/11_-coupled G protein-coupled receptors (GPCRs) related to pleiotropic physiological functions. Upon stimulation by two endogenous neuropeptides, neuromedin U and S (NMU and NMS) with similar binding affinities, NMUR1 and NMUR2 primarily display distinct peripheral tissue and central nervous system (CNS) functions, respectively, due to their distinct tissue distributions. These NMU receptors have triggered extensive attention as drug targets for obesity and immune inflammation. Specifically, selective agonists for NMUR1 in peripheral tissue show promising long-term anti-obesity effects with fewer CNS-related side effects. However, the mechanisms of peptide binding specificity and receptor activation remain elusive due to the lack of NMU receptor structures, which hamper drug design targeting NMU receptors. Here, we report four cryo-electron microscopy structures of G_q_ chimera-coupled NMUR1 and NMUR2 bound with NMU and NMS. These structures present the conserved overall peptide-binding mode and reveal the mechanism of peptide selectivity for specific NMURs, as well as the common activation mechanism of the NMUR subfamily. Together, these findings provide insights into the molecular basis of the peptide recognition selectivity and offer a new opportunity for designing selective drugs targeting NMURs.

## Introduction

Human neuromedin U (NMU) is a 25-amino-acid endogenous peptide that was first discovered in extracts of the porcine spinal cord with a potent smooth muscle contractile activity ^1^. It is also involved in pleiotropic physiological functions, including the regulation of blood pressure, food uptake, nociception, pain perception, bone formation, and immunological responses ^2^. More recently, human neuromedin S (NMS), a 33-amino-acids endogenous peptide, was discovered, which shares an identical C-terminal heptapeptide with NMU. Unlike NMU, which is widely distributed in the central nervous system (CNS) and peripheral tissues, NMS mainly exists in the suprachiasmatic nucleus in the CNS and primarily regulates biological rhythms ^3,4^. Both peptides stimulate two different class A G protein-coupling receptors (GPCRs), neuromedin U receptor 1 (NMUR1) and neuromedin U receptor 2 (NMUR2), with sub-nanomolar affinity but low selectivity ^5,6,7^.

Upon stimulation by NMU and NMS, both NMUR1 and NMUR2 predominantly activate G_q/11_ with some evidence of G_i_ coupling ^8^. The biological functions of the two NMUR subtypes differ by their distinct tissue distributions. NMUR1 is predominantly expressed in peripheral tissues, while NMUR2 is widely distributed in the CNS, most abundantly in the cerebral cortex and hypothalamus ^9^. Both receptor subtypes are closely related to the regulation of food intake and energy balance. Peripheral and central administration of NMU reduced food intake and weight gain by stimulating NMUR1 and NMUR2, respectively ^10-12^. Compared with the NMUR1-selective agonist, the NMUR2 selective agonist has a more potent body weight-lost effect and cause less diarrhea, making it a more well-balanced drug for the treatment of obesity ^13^. Thus, development of selective agonists will benefit from the identification of the mechanisms through which the receptors interact with the peptide ligands.

Extensive efforts have been devoted to understanding the peptide-binding mechanisms of NMUR subtypes. Both NMU and NMS share the highly conserved C-terminal heptapeptide (FLFRPRN-NH2) and the amidated asparagine at the C-terminus across different species ^9,14,15^, indicative of the importance of this conserved peptide segment for receptor recognition. Indeed, this heptapeptide is strongly related to the binding activity, with even single amino acid substitutions reducing their biological effects ^16-18^. Furthermore, the amidated asparagine is also critical for the activity of peptides ^9^. Based on this conserved heptapeptide, a series of NMU analogs have been designed, aiming to develop NMUR1/2 selective agonists. These findings have provided clues for understanding receptor subtype selectivity and designing drug candidates for anti-obesity therapy ^18-25^. Although considerable efforts have been made, the mechanism of peptide recognition by receptors remains to be fully clarified due to the lack of NMUR structures, which has hindered the development of receptor-selective agonists. Here, using single-particle cryo-electron microscopy (cryo-EM), we report four structures of G_q_ chimera-coupled NMUR1 and NMUR2 bound to either NMU or NMS. These structures provide comprehensive insights into the peptide-binding mode and reveal determinants for recognition selectivity of NMUR subtypes by peptides and offer new opportunities for the rational design of selective pharmaceuticals targeting specific NMU receptor subtypes.

## Results

### Overall structures of NMUR1/2 signaling complexes

To facilitate the expression of NMUR1/2 complexes, we introduced a BRIL tag to the N-termini of the wild-type (WT) full-length receptor ^26–28^. The NMUR1-G_q_ chimera complex was stabilized by the NanoBiT strategy ^29^. These modifications have little effect on the pharmacological properties of the NMURs (Supplementary Fig. 1). The Gα_q_ chimera was generated based on the mini-Gα_s_ scaffold with an N-terminus replacement of corresponding sequences of Gα_i1_ to facilitate the binding of scFv16 ^3,31,32^, designated as mGα_s/q/iN_. Unless otherwise stated, G_q_ refers to the mG_s/q/iN_, which was used for structural studies. Incubation of NMU/NMS with membranes from cells co-expressing receptors and heterotrimer G_q_ proteins in the presence of scFv16 ^33–38^ enables effective assembly of NMU/NMS-NMURs-G_q_ complexes, which produces high homogenous complex samples for structural studies.

The structures of the NMU-NMUR1-G_q_-scFv16 and NMS-NMUR1-G_q_-scFv16 complexes were determined by single-particle cryo-EM to the resolutions of 3.2 Å and 2.9 Å, respectively (Fig. 1c, d, Supplementary Fig. 2, and Supplementary Table 1). The cryo-EM structures of NMUR2-G_q_-scFv16 complexes bound to NMU and NMS were determined at 2.8 Å and 3.2 Å, respectively (Fig. 1e, f and Supplementary Fig. 3). The ligand, receptor, and the α5 helix of the Gα_q_ subunit in the four complexes are clearly visible in the EM maps (Supplementary Fig. 4), and side chains of the majority of amino acid residues are well-defined in all components. Hence, these structures provide detailed information on the binding interface between peptides and NMUR1/2, as well as the coupling interface between receptors and G_q_ heterotrimer.

**Fig. 1.**
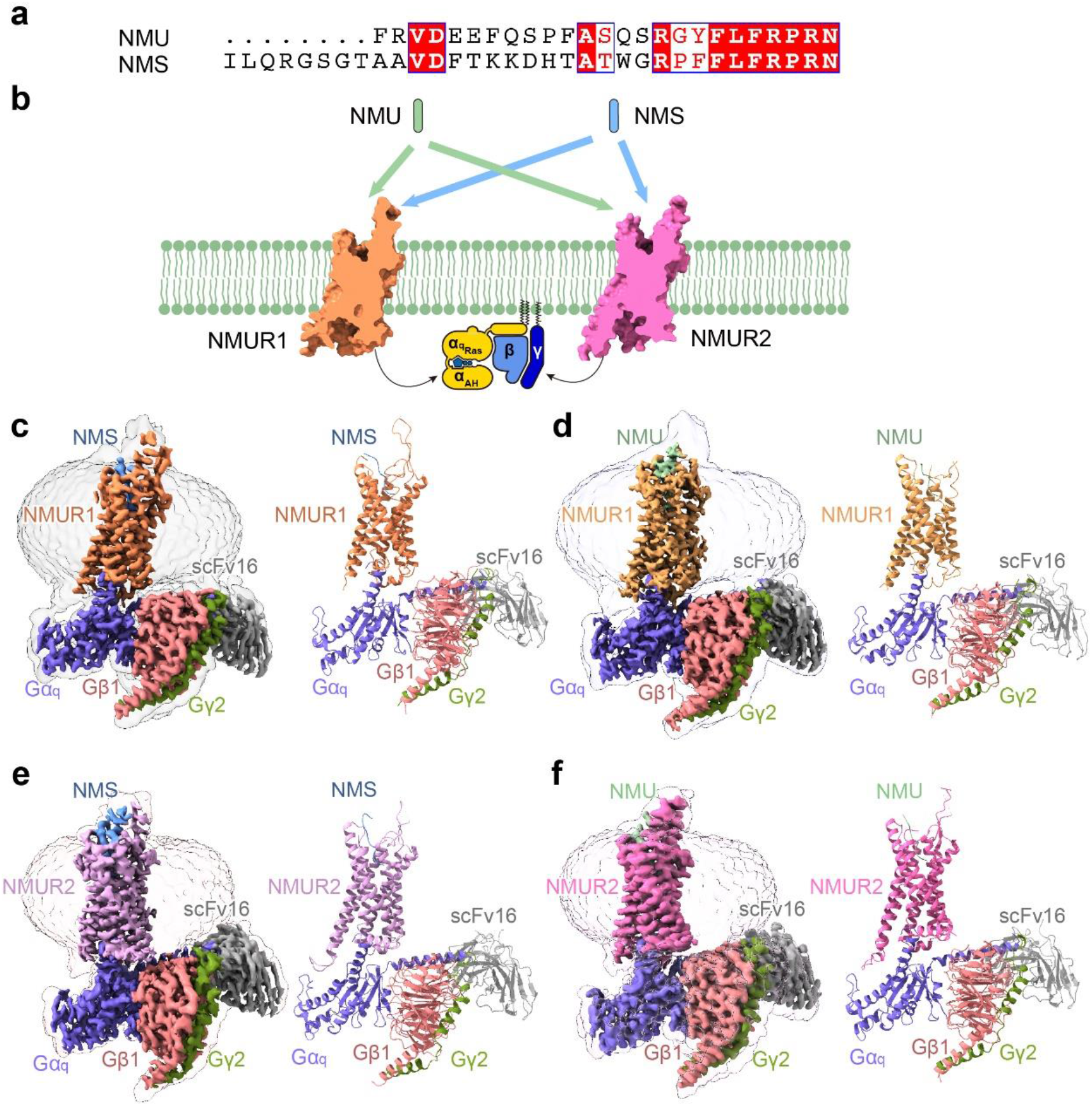
Overall structures of G_q_-coupled NMUR1/2 complexes bound to NMU and NMS. **a** Sequence alignment of NMU and NMS created by CLUSTALW (https://www.genome.jp/tools-bin/clustalw) and ESPript 3.0 (https://espript.ibcp.fr/ESPript/cgi-bin/ESPript.cgi). **b** Schematic illustration of peptide-binding and G_q_ protein-coupling of NMURs. **c-f** Orthogonal views of the density maps and models of NMU-NMUR1-G_q_-scFv16 (**c**), NMS-NMUR1-G_q_-scFv16 (**d**), NMU-NMUR2-G_q_-scFv16 (**e**), and NMS-NMUR2-G_q_-scFv16 (**f**) complexes. NMS is shown in light blue, NMS-bound NMUR1 in orange, and NMS-bound NMUR2 in plum. NMU is displayed in green, NMU-bound NMUR1 in brown, and NMU-bound NMUR2 in hot pink. The G_q_ heterotrimer is colored by subunits. Gα_q_, purple; Gβ1, salmon; Gγ2, dark green; scFv16, grey. Gα_q_ refers to mGα_s/q/iN_.

The overall conformations of the four active NMUR1/2-G_q_ complexes are highly similar (Fig. 1c-f and Supplementary Fig. 5a), with root mean square deviation (R.M.S.D.) values of 0.371-0.735 Å for the entire complexes and 0.467-0.794 Å for the receptor. Unlike most GPCRs with a solved structure, the EM density of extracellular loop 2 (ECL2) from both receptors is oriented almost parallel to the transmembrane domains (TMDs). Interestingly, ambiguous EM densities of the N-termini of peptides can be observed in these four complexes. These N-termini of the peptides seem to interact with the ECL2s, consistent with the previous report that ECL2 is involved in peptide-induced receptor activation (Supplementary Fig. 5b) ^39,40^. In addition, the binding poses of NMU and NMS in both receptors are highly overlayed (R.M.S.D. of 0.749 Å for NMUR1 and 0.527 Å for NMUR2). Although occupying the same TM cavity for all peptide GPCR structures determined to date ^41,42,43^, NMU and NMS adopt different binding poses, demonstrating the diverse recognition modes of peptides (Fig. 2a-f and Supplementary Fig. 5c).

**Fig. 2.**
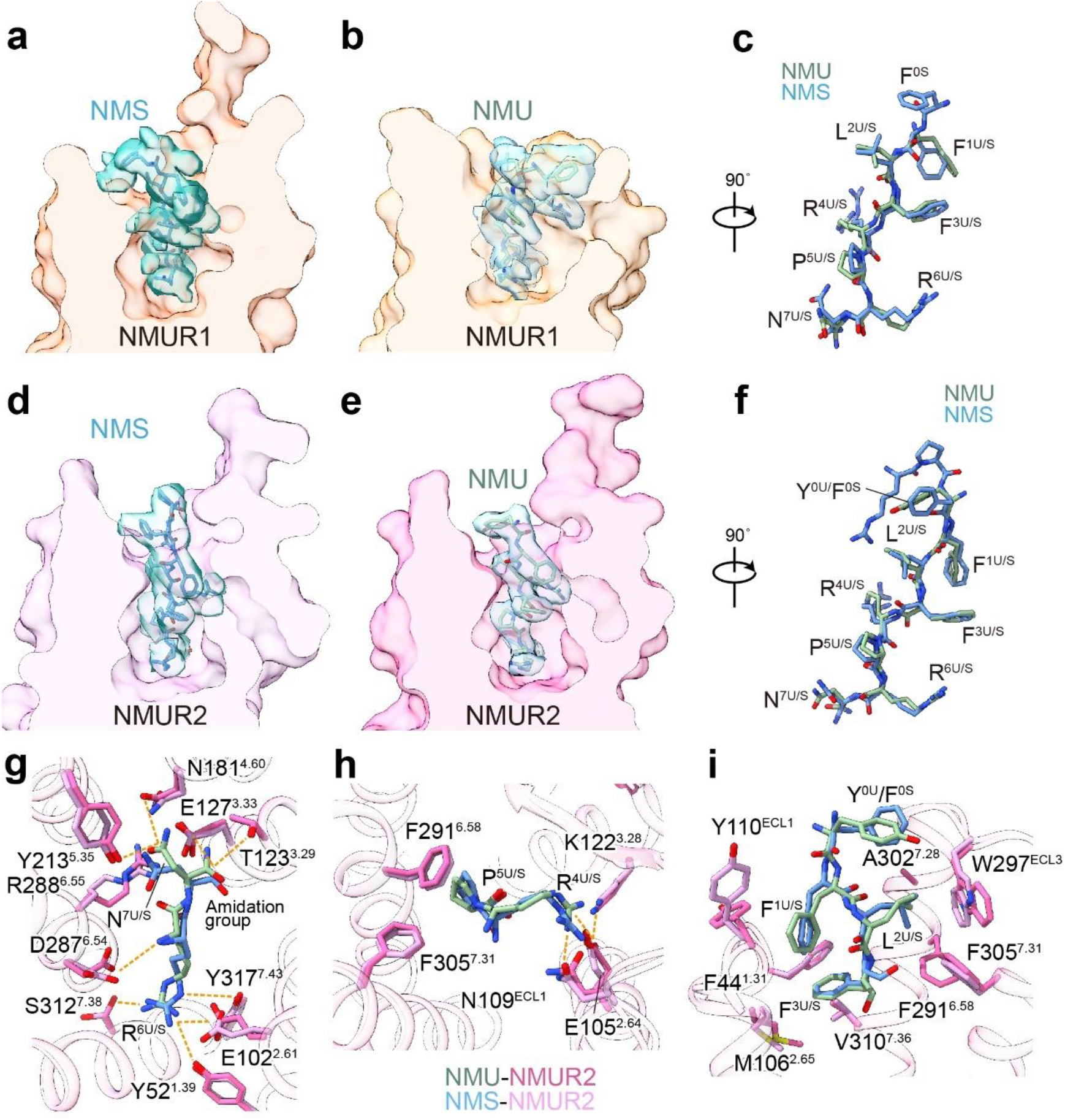
The conserved binding pocket of NMUR2. **a-c** Cut-away view of NMU/NMS binding pocket of NMUR1 (**a, b**) and structural superposition of NMU and NMS in NMUR1 (**c**). **d-f** Cut-away view of NMU/NMS binding pocket of NMUR2 (**d, e**) and structural superposition of NMU and NMS in NMUR1 (**f**). The density of NMS is colored in dark green, while the density of NMU is colored in blue. **g-i** Detailed interaction of NMU/NMS with residues in NMUR2. The binding site of N^7U/S^ and R^6U/S^ (**g**), P^5U/S^ and R^4U/S^ (**h**), F^3U/S^, L^2U/S^, F^1U/S^, and Y^0U^/F^0U^ (**i**) are shown. Hydrogen bonds and salt bridges are depicted as orange dashed lines. NMU and NMS are shown as sticks. NMS is shown in light blue and NMS-bound NMUR2 in plum. NMU is displayed in green and NMU-bound NMUR2 in hot pink.

### Binding modes of NMU and NMS for NMURs

NMU and NMS in the four NMURs complex structures adopt similar conformations. C-termini of both peptides insert into an overlapped orthosteric binding pocket, comprising all TM helices and ECLs (Fig. 2a-f, Supplementary Figs. 6 and 7). Due to the sequence consensus of the C-terminal heptapeptide, both NMU and NMS share highly conserved binding modes for specific NMUR subtypes. We use the structure of the NMU-NMUR2-G_q_ complex, which shows a higher resolution relative to the NMS-bound one, to analyze the peptide binding mode for NMUR2.

At the bottom region of the orthosteric peptide-binding pocket, polar receptor residues form an extensive polar interaction network with R^6U^ and amidated N^7U^ (Fig. 2g). The amidation group of N^7U^ makes a polar contact with E127^3.33^, structurally supporting the fact that this amidation modification is necessary for the activity of NMU ^44^. The side chain of N^7U^ forms H-bond interactions with N181^4.60^, Y213^5.35^, and R288^6.55^. Noteworthily, a conserved salt bridge between E127^3.33^ and R288^6.55^ exists in NMURs and other GPCRs with relatively high homology, including ghrelin and neurotensin receptors (Supplementary Fig. 5d). This conserved salt bridge may closely pack TM3 and TM6, thereby preventing peptides from further insertion and stabilizing the active receptor conformation. On the opposite orientation of N^7U^, R^6U^ was fastened mainly through polar interactions by S3I2^7.38^ and E102^2.61^, the latter further making intramolecular polar contacts with Y52^1.39^ and Y317^7.43^. These extensive polar interaction networks mediated by R^6U^ and amidated N^7U^ make substantial contributions to NMU activity, which is supported by the alanine mutagenesis analysis (Fig. 2g, Supplementary Figs. 8 and 9, Supplementary Table 2). Another polar network links R^4U^ to E105^2.64^, N109^ECL1^, and K122^3.28^, locking NMU with TM2, TM3, and ECL1 (Fig. 2h). Apart from the polar interaction networks, F^1U^, L^2U^, and F^3U^ are engaged in hydrophobic contacts with the upper part of the TMD pocket. F^1U^ and F^3U^ form intramolecular π-stacking and hydrophobically interact with the F44^1.31^, M106^2.65^, Y110^ECL1^, and V310^7.36^ (Fig. 2i). L^2U^ faces an environment composed of F291^6.58^, W297^ECL3^, A302^7.28^, and F305^7.31^. F291^6.58^ and F305^7.31^ are also involved in the hydrophobic interactions with P^4U^, extending the L^2U^-mediated hydrophobic interaction network (Fig. 2h, i). Most of these hydrophobic residues are involved in NMU-induced NMUR2 activation. It should be noted that the hampered peptide activities on Y110^ECL1^A and K122^3.28^A mutants are probably be attributed to the decreased expression level (Supplementary Figs. 8 and 9, Supplementary Table 2). The C-terminal heptapeptide and the amidated asparagine of NMS (F^1S^-N^7S^-NH_2_) share a highly similar binding mode with NMU for NMUR2 (Fig. 2g-i). This conserved peptide-binding pattern is also observed in NMUR1 (Supplementary Fig. 6).

Noteworthily, cognate residues for both NMUR1 and NMUR2 surrounding P^5U/S^-N^7U/S^, the three amino acids at the end of the peptides, are completely conserved, thus making highly similar interactions with the C-termini of peptides. In contrast, the peptide segment F^1U/S^-R^4U/S^ of both peptides face distinct physicochemical environments and differ in the interaction pattern for the two NMUR subtypes (Fig. 3 and Supplementary Figs. 6). This distinction of the F^1U/S^-R^4U/S^ binding environment may provide a basis for discriminating selective agonists by specific NMUR subtypes, thus probably offering an opportunity for designing NMUR subtype-selective ligands.

**Fig.3.**
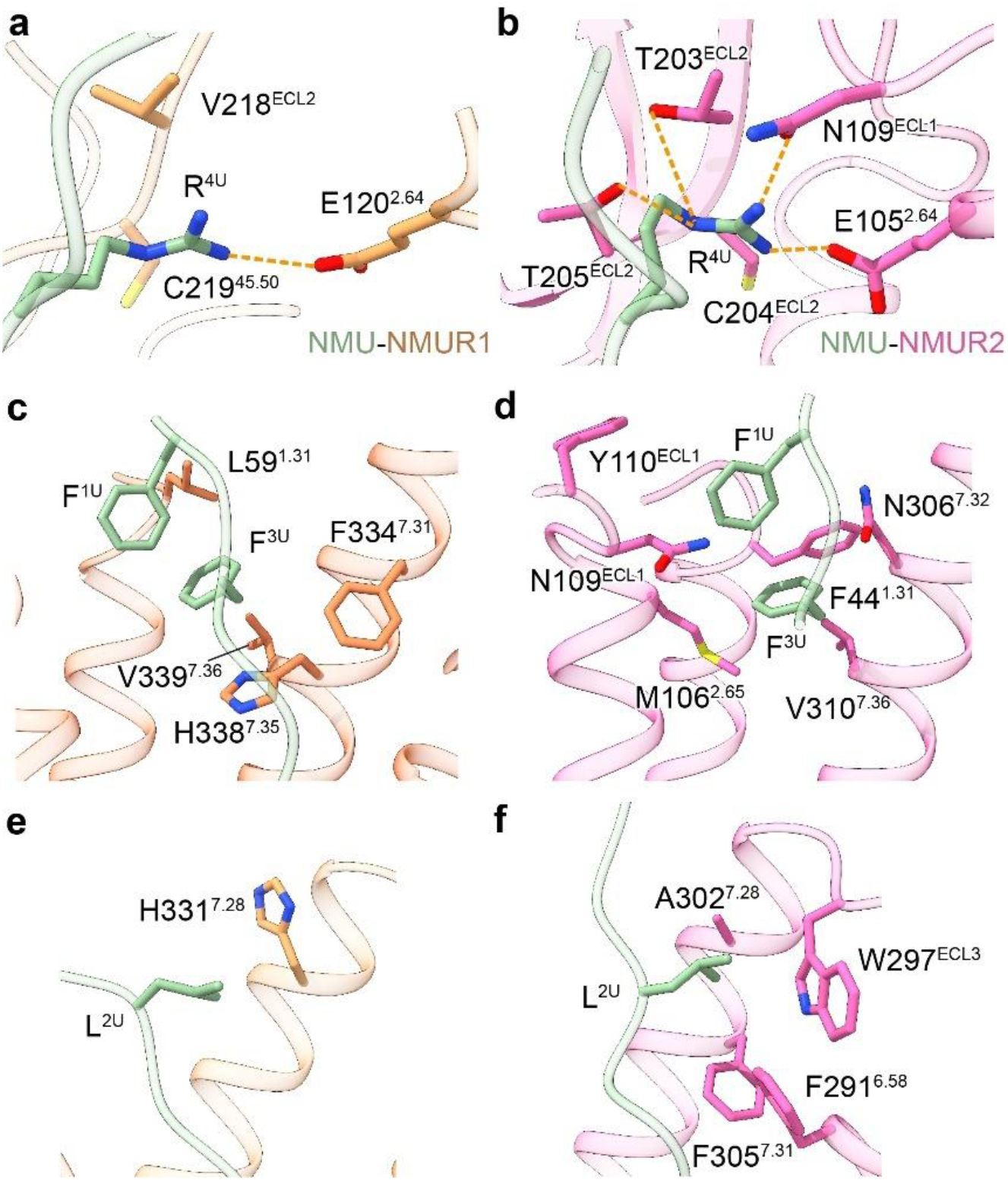
Comparison of the binding mode L^2^-F^3^-R^4^ in NMU between NMUR1 and NMUR2. Detailed interactions between R^4U^ (**a, b**), F^3U^ (**c, d**), and L^2U^ (**e, f**) and pocket residues in NMUR1 and NMUR2 are shown. Side chains of residues are displayed in sticks. Hydrogen bonds and salt bridges are depicted as orange dashed lines. NMU is displayed in green, NMUR1 in brown, and NMUR2 in hot pink.

### Molecular basis of peptide selectivity for NMURs

Hexapeptide analogs of NMU with amino acid substitution at L^2U^-F^3U^-R^4U^ have shown potential selectivity for specific NMUR subtypes ^18,19,21,22^. Pairwise structures of NMURs in complex with NMU offer a template for understanding the selective recognition basis of these NMU analogs.

For NMUR2, R^4U^ lies in a more potent polar environment (E105^2.64^, N109^ECL1^, T203^ECL2^, and T205^ECL2^) than NMUR1 (E120^2.64^). Moreover, the side chain of R^4U^ in NMUR1 is less stretched due to the steric hindrance caused by ambient residues (Fig. 3a, b). Replacing the side chain of R^4U^ with the aminoalkyl group with a comparable or shorter carbon chain decreased their activity to NMUR1 ^18^. However, interactions between these substituted side chains and residues in NMUR2 are more easily maintained, providing the NMUR2 binding preference of these NMU analogs. Similarly, guanidine derivatives with shorter carbon chains also displayed higher selectivity for NMUR2 over NMUR1 ^18,25^. According to the molecular docking results, the guanidinium group may polarly interact with T203^ECL2^ and T205^ECL2^ in NMUR2 but fail to engage with cognate hydrophobic residues in NMUR1 (V218^ECL2^ and C219^ECL2^) (Supplementary Fig. 10a-c). On the contrary, guanidine and aminoalkyl derivatives with comparable or longer carbon chains relative to arginine showed non-selectivity or slightly increased selectivity to NMUR1 ^18^.

In contrast to NMUR2, a more extensive hydrophobic network surrounding F^3U^ in NMUR1 (L59^1.31^, F334^7.31^, H338^7.35^, and V339^7.36^) probably makes a greater contribution to stabilizing peptide-NMUR1 interaction, thus raising a hypothesis that this hydrophobic network may discriminate peptide derivatives with different receptor selectivity (Fig. 3c, d). This hypothesis is supported by the fact that substituting the aromatic phenyl ring of F^3U^ by an aliphatic cyclohexyl ring or other alkyl side chains with weaker hydrophobicity increased their binding preference for NMUR2 ^18,19^. An isopropyl and cyclohexyl substitution of the F^3U^ side chain may maintain hydrophobic interactions with M106^2.65^ and V310^7.36^ in NMUR2, which are absent in NMUR1 (Supplementary Fig. 10d, e). Conversely, displacing the side chain of F^3U^ with a biphenyl, naphthyl, or indolyl group enhanced the binding selectivity for NMUR1 by forming hydrophobic interactions with L59^1.31^, F334^7.31^, and H338^7.35^ ^19,21^, thus probably maintaining or even enhancing its interaction with NMUR1. In contrast, steric hindrance may occur between bulky side-chain substitution and residues in NMUR2, limiting its binding to NMUR2 (Supplementary Fig. 10f-h).

For NMUR2, L^2U^ was buried in a compact residue environment (F291^6.58^, W297^ECL3^, A302^7.28^, and F305^7.31^), meaning that it is unable to accommodate bulky side-chains. Conversely, a wider space surrounding L^2^ in the NMUR1 pocket may serve as a determinant for designing NMUR1-selective agonists (Fig. 3e-f). Indeed, the heteroaromatic ring and bulky aromatic ring substitution of the L^2U^ side-chain are crucial to developing an NMUR1-selective agonist ^18,21,23^. Our molecular docking analysis reveals that a biphenyl, naphthyl, or indolyl substitution of L^2U^ side-chain sits closer to H331^7.28^ and may create extra interactions with NMUR1 relative to NMUR2 (Supplementary Fig. 10i-k). It should be noted that although the connectivity of our structural observation and the previous functional evidence on peptide selectivity, we cannot completely exclude the possible impact of the NanoBiT, which is introduced in the structure determination of NMUR1 complexes. Together, combined with previous functional findings, our structures enhance our understanding on the basis of NMUR subtype selectivity and offer a template for designing agonists targeting specific NMUR subtypes.

### Activation mechanism of NMURs

Since the complexes of NMUR1 and NMUR2 with NMU or NMS share highly overlaid overall conformations, we applied the structure of the NMU-NMUR2-G_q_ complex to consider the activation mechanisms of NMURs. Structural comparison of this complex with the antagonist-bound ghrelin receptor supports the contention that these NMURs are in the active state, featured by the pronounced outward displacement of the cytoplasmic end of TM6 and concomitantly inward shift of TM7 (Fig. 4a).

**Fig. 4.**
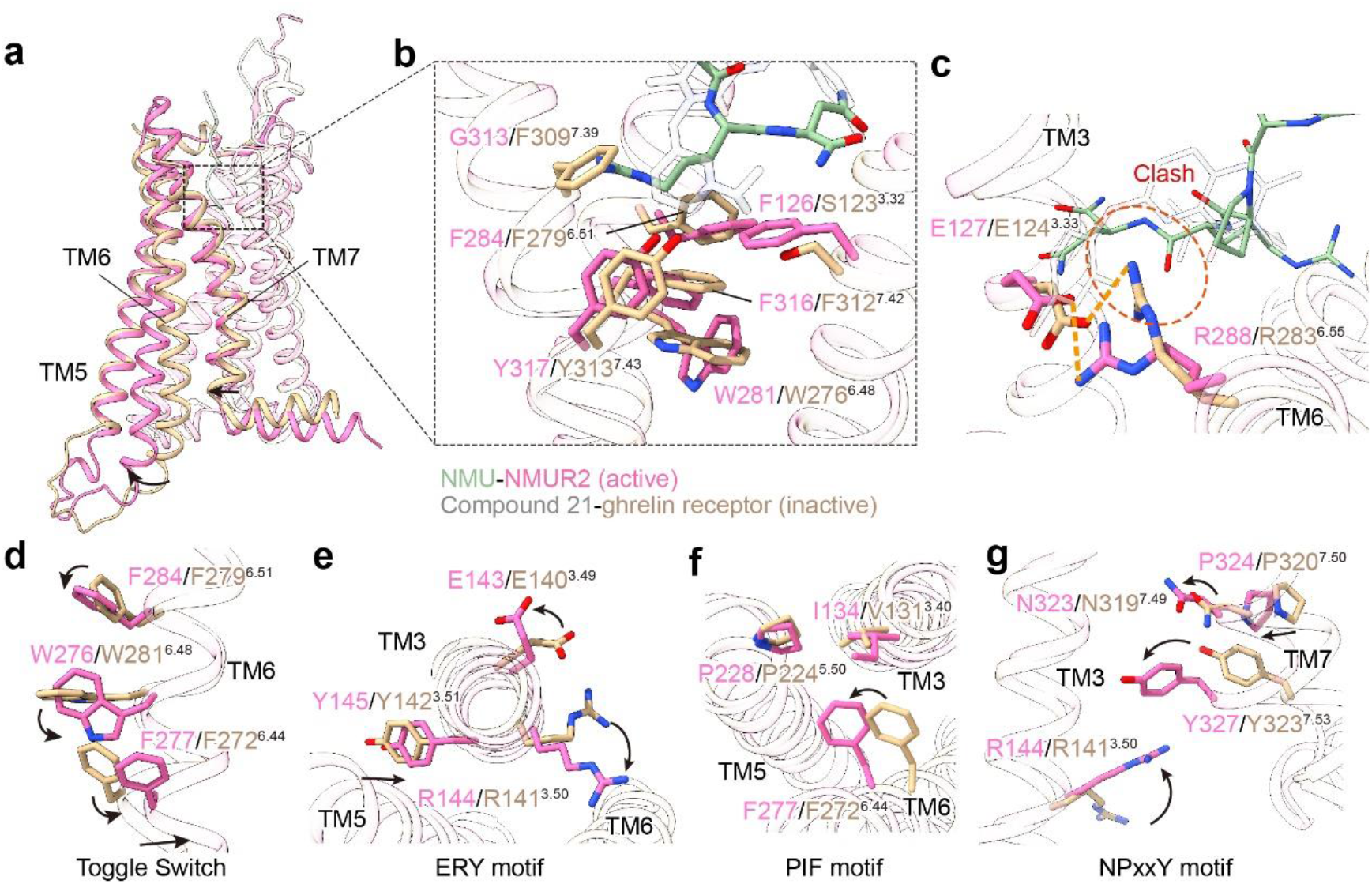
Activation mechanism of NMURs. **a** Structural superposition of active NMUR2 and antagonist-bound ghrelin receptor (PDB 6KO5) from the side view. The movement directions of TMs in NMUR2 relative to the ghrelin receptor are highlighted as black arrows. NMUR2 and ghrelin receptors are colored in hot pink and wheat, respectively. **b** Interactions between NMU and residues located at the bottom of peptide binding pocket. **c** Comparison of interaction between peptide and R^6.55^. The possible clash is highlighted by a red dashed circle. Hydrogen bonds and salt bridges are depicted as orange dashed lines. Compound 21, the antagonist of ghrelin receptor, and NMU are colored in grey and green, respectively. **d-g** Conformational changes of the conserved “micro-switches” upon receptor activation, including Toggle switch (**d**), ERY (**e**), PIF (**f**), and NPxxY (**g**) motifs. The conformational changes of residue side chains are shown as black arrows upon NMUR2 activation in contrast to the antagonist-bound ghrelin receptor.

Due to the steric hindrance caused by a hydrophobic lock comprising of F284^6.51^, F126^3.32^, and Y317^7.43^, NMU is not able to directly contact the “toggle switch” residue W281^6.48^, which often undergoes a movement upon ligand binding ^45,46^ (Fig. 4b). Alternatively, the side chain of the amidated N^7U^ in NMU may push the side chain of R288^6.55^, causing it to swing away from the receptor helical core (Fig. 4c). Concomitantly, the swing of R288^6.55^ may lead to the conformational changes of F284^6.51^ and W281^6.48^, further leading to the swing of F277^6.44^ and the pronounced outward displacement of the cytoplasmic end of TM6 (Fig. 4d). The other conserved residues in “micro-switches” (ERY, PIF, and NPxxY) also undergo active-like conformational changes relative to the antagonist-bound ghrelin receptor and transmit the peptidic agonism signaling to the cytoplasmic face of the receptor to facilitate G protein coupling (Fig. 4e-g). Also, rotameric switches were caused by the conformational changes of F284^6.51^ and W281^6.48^. The repacking of the inter-helical hydrophobic contacts between TM6 and TM7 occurred that led to the inward shift of the cytoplasmic end of TM7 (Fig. 4a). The R^6.55^-mediated activation mechanism shared by NMUR1 is also captured in the ghrelin receptor ^31^ and neurotensin receptor 1 ^47^, probably serving as a common mechanism across other peptide GPCRs with high sequence homology with NMURs, including the motilin receptor and neurotensin receptor 2 (Supplementary Fig. 5d).

### The interface between NMURs and the Gα_q_ subunit

The structure of the NMU-NMUR2-G_q_ complex was applied to characterize the interface between NMURs and G_q_ heterotrimer in the detergent micellular environment. Like other G protein-coupled GPCRs, the primary NMUR2-Gα_q_ subunit interface is comprised of the C-terminal helix (α5 helix) of Gα_q_ and the cytoplasmic cavity of the TMD core (Fig. 5a). Structural comparisons of NMUR2-G_q_ with G_q_-coupled cholecystokinin A receptor (CCK_A_R, PDB 7EZM ^48^), histamine H1 receptor (H_1_R, PDB 7DFL ^49^), and G_11_-coupled muscarinic acetylcholine receptor M1 (M_1_R, PDB 6OIJ ^50^) complexes reveal distinct NMURs-G_q_ coupling features. The NMU-NMUR2-G_q_ complex displays a similar overall conformation with the CCK_A_R-G_q_ complex but differs in conformations of TM6 and the Gα subunit relative to M_1_R-G_11_ and H_1_R-G_q_ complexes. Compared with G_q/11_-coupled M_1_R and H_1_R, the TM6 of NMUR2 undergoes a remarkably inward displacement (Fig. 5a). Consequently, the extreme C-terminal α5 helix of Gα_q_ subunit in NMUR2-G_q_ complex shifts inward toward TM2, TM3, and ICL2 to avoid clashes with TM6, accompanied with the rotation of the entire Gα_q_ subunit (Fig. 5a, b). Specifically, in contrast to Y356^H5.23^ (measured at Cα atom of L^H5.25^, superscript refers to CGN system^51^) in the M_1_R-G_11_ complex, the hydroxyl of Y358 ^H5.23^ shift ~4 Å to create additional interactions with TM2 and ICL2 (T80^2.39^ and S158^ICL2^) of NMUR2. Similar interactions were observed between Y356 ^H5.23^ and T76^2.39^ and Q153^ICL2^ of CCK_A_R (Fig. 5b, c). On the contrary, Y356 ^H5.23^ is anchored by polar interactions with S126/S128^3.53^ and R137/R139^ICL2^ in M_1_R/H_1_R (Fig. 5d). Together, these findings reveal the specific nature of the NMURs-G_q_ interface. These NMURs-G_q_ complex structures are added to the pool for enhancing the understanding of the GPCR-G_q_ coupling mechanism.

**Fig. 5.**
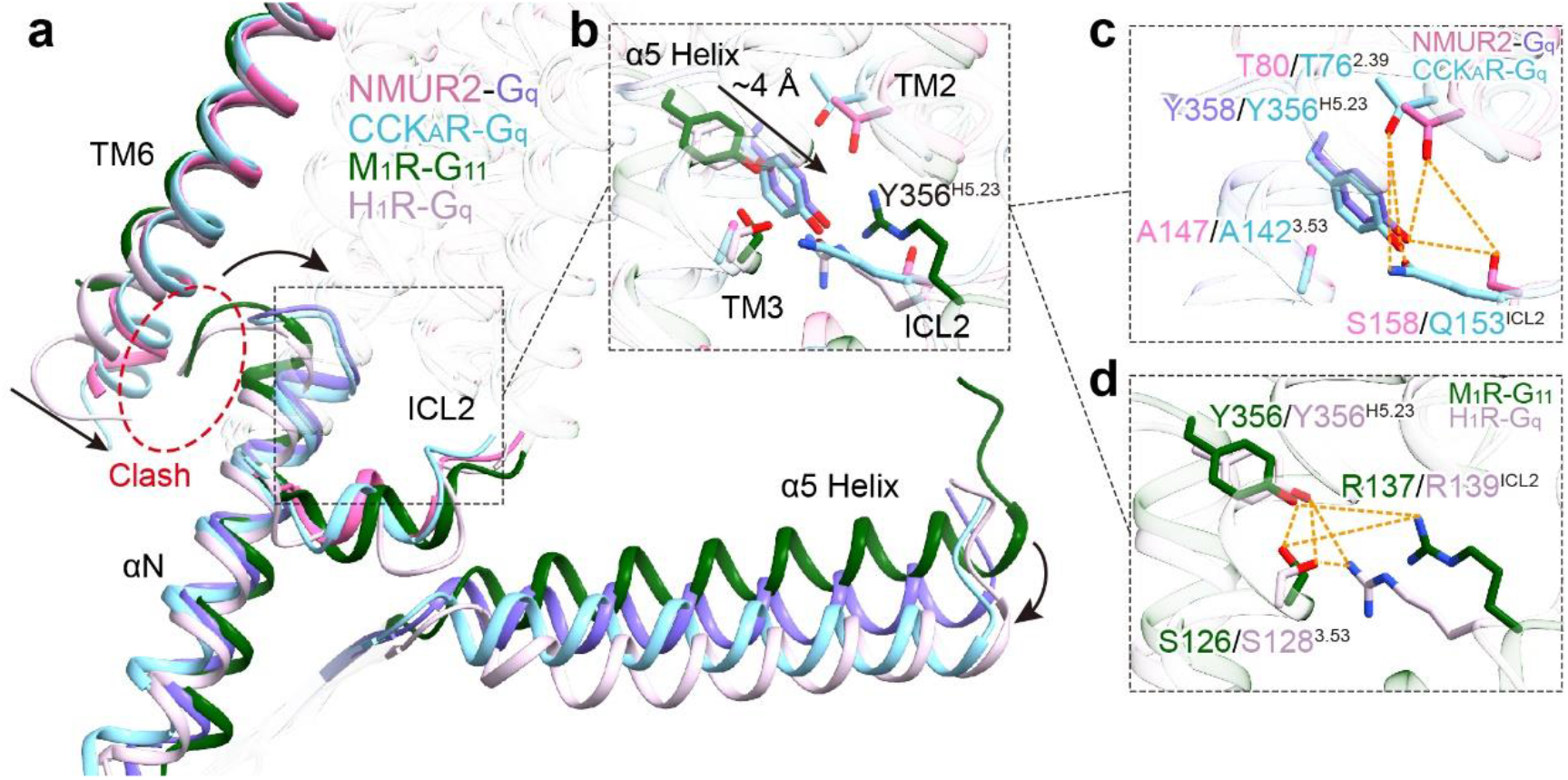
G_q_ protein-coupling of NMUR2. **a** An overall conformational comparison of G_q_-coupled NMUR2 with G_q_-coupled CCK_A_R (PDB 7EZM), H_1_R (PDB 7DFL), and G_11_-couple M_1_R (PDB 6OIJ). TM6 and ICL2 of receptors, as well as αN and α5 helix of G proteins, are highlighted. The potential clashes between TM6 of receptors and α5 helices of Gα_q/11_ subunits are highlighted by dashed circles. **b-d** Interactions between Y^H5.23^ of Gα_q/11_ α5 helices and receptors, including NMUR2, M_1_R, CCK_A_R, and H_1_R. The detailed polar interactions between Y^H5.23^ and NMUR2 and CCK_A_R (**c**), as well as M_1_R and H_1_R (**d**) are shown. The polar interactions are shown by orange dashed lines. The displacements of components in G_q_-coupled NMUR2 and CCK_A_R relative to G_q/11_-coupled M_1_R and H_1_R are indicated by black arrows. Colors are shown as indicated. G_q_ refers to G_q_ chimeras used in structural studies of the four GPCRs. G_q_ coupled by NMUR2 refers to mG_s/q/iN_.

## Discussion

In this paper, we reported four cryo-EM structures of G_q_-coupled NMUR1 and NMUR2 bound to either NMU or NMS. These structures present a conserved orthosteric peptide-binding pocket in both NMUR subtypes, which accommodate the identical heptapeptide at the C-termini of NMU and NMS. Combining structural observation and alanine mutagenesis analysis reveals the binding mode of the C-terminal heptapeptide, which is critical for the activity of both peptides. Intriguingly, we observed an ambiguous EM density in proximity to ECL2 in the map of our complexes except for the NMU-NMUR1-G_q_ complex, which is derived from the N-terminus of NMU and NMS with high probability. This observation indicates a direct contact between the N-terminal segment of peptides and ECL2, consistent with the previous report that the N-termini of peptides made a substantial contribution to its binding activity to NMURs ^6,39,40,52^. Moreover, pairwise structural comparison of NMUR1 and NMUR2 reveals potential determinants for receptor subtype selectivity. Additionally, a mechanism of R^6.55^-triggered receptor activation was found, which is conserved by the ghrelin receptor and neurotensin receptor 1^31,47^.

These structures provide a template for understanding the mechanism underlying peptide recognition selectivity for NMURs and offer an opportunity for designing receptor-selective ligands (Supplementary Fig. 11). The extreme C-terminal tripeptide with amidated modification (P^5^-R^6^-N^7^-NH2) is buried in a potent polar binding pocket, which is highly conserved between the two NMUR subtypes. In contrast, distinct physiochemical environments surrounding a tripeptide (L^2^-F^3^-R^4^) between two NMUR subtypes serve as determinants for NMUR subtype preference. Specifically, substituting R^4^ with a shorter or a weaker polar side chain may maintain the original polar interactions with NMUR2 relative to NMUR1, thus enhancing the NMUR2 selectivity. The side-chain substitution of F^3^ displays a double-edged role in both NMUR1 and NMUR2 selectivity. Displacing the aromatic ring of F^3^ with a smaller hydrophobic or a less aromatic side-chain improves NMUR2 selectivity. On the contrary, F^3^ bearing a bulkier hydrophobic substituent enhances NMUR1 selectivity. Additionally, a peptide analog bearing bulky groups relative to L^2^ may maximize its abundant space and avoid the steric hindrance, thus delivering a higher selectivity on NMUR1. Single or combined substitutions of L^2^-F^3^-R^4^ side chains may provide novel drug candidates with NMUR subtype selectivity for anti-obesity therapy.

## Author Contributions

C.Y. screened the expression constructs, optimized the NMURs-G_q_ complexes, prepared the protein samples for final structure determination, participated in cryo-EM grid inspection, data collection, and model building; C.Y. and Y.Z designed the mutations and executed the functional studies; C.Y., P.X., and S.H. build and refine the structure models; W.Y. designed G_q_ protein constructs; H.E.X. and Y.J. conceived and supervised the project; C.Y. and Y.J. prepared the figures and drafted manuscript; Y.J. wrote the manuscript with inputs from all authors.

## Method

### Constructs

The full-length human NMUR1 and NMUR2 were modified to contain the N-terminal thermally stabilized BRIL ^26^ to enhance receptor expression and the addition of affinity tags, including an N-terminal Flag tag and a 10×His-tag. LgBiT was inserted at the C-terminus of the human NMUR1 using homologous recombination. Both modified NMUR1 and NMUR2 were cloned into the pFastBac (Thermo Fisher Scientific) vectors using the ClonExpress II One Step Cloning Kit (Vazyme Biotech). An engineered Gα_q_ chimera was generated based on the mini-Gα_s_ scaffold with its N-terminus replaced by corresponding sequences of Gα_i1_, designated as mGα_s/q/iN_. Human wild-type (WT) Gβ1, human Gγ2, and a single-chain antibody scFv16 ^53^, as well as a Gβ1 fused with SmBiT at its C-terminus, were cloned into pFastBac vectors.

### Insect cell expression

Human NMUR1, NMUR2, G_q_ chimera, Gβ1, Gγ, scFv16, and Ric8a were co-expressed in High Five insect cells (Invitrogen) using the baculovirus method (Expression Systems). Cell cultures were grown in ESF 921 serum-free medium (Expression Systems) to a density of 2-3 million cells per mL and then infected with six separate baculoviruses at a suitable ratio. The culture was collected by centrifugation 48 h after infection, and cell pellets were stored at −80°C.

### Complex purification

Cell pellets were thawed in 20 mM HEPES pH 7.4, 50 mM NaCl, 10 mM MgCl_2_, and CaCl_2_ supplemented with Protease Inhibitor Cocktail (TargetMol). For the NMU-NMUR1/2-G_q_-scFv16 complexes, 10 μM NMU (GenScript) and 25 mU ml^-1^ apyrase (Sigma) were added. For the NMS-NMUR1/2-G_q_-scFv16 complexes, 5 μM NMS (GenScript) and 25 mU ml^-1^ apyrase (Sigma) were added. The suspension was incubated for 1 h at room temperature, and the complex was solubilized from the membrane using 0.5% (w/v) lauryl maltose neopentylglycol (LMNG) (Anatrace) and 0.1% (w/v) cholesteryl hemisuccinate (CHS) (Anatrace) for 2 h at 4°C. Insoluble material was removed by centrifugation at 65,000 g for 35 min, and the the supernatant was purified by nickel affinity chromatography (Ni Smart Beads 6FF, SMART Lifesciences). The resin was then packed and washed with 20 column volumes of 20 mM HEPES pH 7.4, 50 mM NaCl, 0.01% (w/v) LMNG, and 0.002% CHS. The complex sample was eluted in buffer containing 300 mM imidazole and concentrated using an Amicon Ultra Centrifugal Filter (MWCO 100 kDa). The complex was then subjected to size-exclusion chromatography on a Superdex 200 Increase 10/300 column (GE Healthcare) pre-equilibrated with size buffer containing 20 mM HEPES pH 7.4, 100 mM NaCl, 0.00075% (w/v) LMNG, 0.00025% (w/v) GDN (Anatrace) and 0.00015% CHS to separate complexes. For the NMU-bound or NMS-bound complexes, 10 μM NMU and 5 μM NMS were included in the Size Buffer, respectively. Eluted fractions were evaluated by SDS-PAGE and those consisting of receptor-G_q_ protein complex were pooled and concentrated for cryo-EM experiments.

### Cryo-EM grid preparation and data acquisition

Three microliters of the purified NMUR1 and NMUR2 complexes at around 18 mg ml^-1^, 15 mg ml^-1^, 20 mg ml^-1^, and 15 mg ml^-1^ for NMU-NMUR1, NMS-NMUR1, NMU-NMUR2, and NMS-NMUR2 complexes, respectively, were applied onto a glow-discharged Quantifoil R1.2/1.3 200-mesh gold holey carbon grid. The grids were blotted for 3 s under 100% humidity at 4°C and then vitrified by plunging into liquid ethane using a Vitrobot Mark IV (Thermo Fisher Scientific). For all NMURs complexes, Cryo-EM data collection was performed on a Titan Krios G3i at a 300 kV accelerating voltage at the Shuimu BioSciences Ltd (Beijing, China) and the micrographs were recorded using a super-resolution counting mode at a pixel size of 0.54 Å. Micrographs were obtained at a dose rate of about 18.5 e Å^-2^ s^-1^ with a defocus ranging from −1.0 to −3.0 μm. Each micrograph was dose-fractionated to 32 frames with a total exposure time of 3.33 s. A total of 3746, 3424, 2993, and 2862 movies were collected for NMU-NMUR1, NMS-NMUR1, NMU-NMUR2, and NMS-NMUR2 complexes, respectively.

### Image processing and 3D reconstruction

Image stacks were subjected to beam-induced motion correction using MotionCor2.1 ^54^. Contrast transfer function (CTF) parameters for each-non-dose-weighted micrograph were determined by Gctf ^55^. Automated particle selection and data processing were performed using RELION-3.0 beta2 ^56^. For the dataset of the NMS-NMUR2-G_q_ complex, particles selection yielded 5,191,427 particles, which were subjected to reference-free 2D classification. The map of the 5-HT_1E_-G_i_ complex (EMD-30975) low-pass-filtered to 30 Å was used as an initial reference model for 3D classification. A further two rounds of 3D classifications focusing the alignment on the complex, except AHD of the Gα subunit, produced one high-quality subset accounting for 728,263 particles. These particles were subsequently subjected to Bayesian polishing, CTF refinement, and 3D refinement, which generated a map with an indicated global resolution of 3.2 Å at a Fourier shell correlation (FSC) of 0.143. Local resolution was determined using the Resmap package with half map as input maps.

For the dataset of NMU-NMUR2-G_q_ complex, particles selection yielded 4,738,667 particles, which were subjected to reference-free 2D classification. The map of the NMS-NMUR2-G_q_ complex low-pass-filtered to 60 Å was used as an initial reference model for 3D classification. A further two rounds of 3D classifications focusing the alignment on the complex, except AHD of the Gα, produced three high-quality subsets accounting for 2,087,642 particles. These particles were subsequently subjected to Bayesian polishing, CTF refinement, and 3D refinement, which generated a map with an indicated global resolution of 2.8 Å at an FSC of 0.143.

For the dataset of the NMU-NMUR1-G_q_ complex, automated particle selection yielded 5,129,300 particles. The particles were extracted on a binned dataset with a pixel size of 1.08 Å and were subjected to a reference-free 2D classification. The map of the NMS-NMUR2-G_q_ complex solved in this study was used as an initial reference model for 3D classification. Further 3D classifications focusing the alignment on the complex, except the α helical domain of the Gα, produced the high-quality subset accounting for 312,310 particles. These particles were subsequently subjected to Bayesian polishing, CTF refinement, and 3D refinement, which generated a map with an indicated global resolution of 3.2 Å at an FSC of 0.143.

For the dataset of the NMS-NMUR1-G_q_ complex, particles selection yielded 4,708,785 particles, which were subjected to reference-free 2D classification. The map of NMU-NMUR1-G_q_ complex low-pass-filtered to 60 Å was used as an initial reference model for 3D classification. A further two rounds of 3D classifications focusing the alignment on the complex, except AHD of the Gα subunit, produced one high-quality subset accounting for 588,662 particles. These particles were subsequently subjected to Bayesian polishing, CTF refinement, and 3D refinement, which generated a map with an indicated global resolution of 2.9 Å at an FSC of 0.143.

### Structure determination and refinement

The cryo-EM structure of the NMS-NMUR2-G_q_ complex was solved using 5-HT_1E_ as the initial model (PDB 7E33). All other three structures of NMURs-G_q_ complexes were built using the NMS-NMUR2-G_q_ model as a template. The models were docked into cryo-EM density maps using Chimera ^57^, followed by iterative manual adjustment and rebuilding in Coot ^58^ and ISOLDE ^59^, against the cryo-EM electron density maps. Realspace and reciprocal refinements were performed using PHENIX ^60^, as well as the model statistics validation. Structural figures were prepared in Chimera ^57^, ChimeraX ^61^, and PyMOL (https://pymol.org/2/). The final refinement statistics are provided in Supplementary Table 1.

### Inositol phosphate accumulation assay

IP-One production was measured using the IP-One HTRF kit (Cisbio) ^62^. Briefly, AD293 cells (Agilent) were grown to a density of 400,000-500,000 cells per mL and then infected with separate plasmids at a suitable concentration. The culture was collected by centrifugation 24 h after incubation at 37°C in 5% CO_2_ with a Stimulation Buffer. The cell suspension was then dispensed in a white 384-well plate at a volume of 7 μl per well before adding 7 μl of ligands. The mixture was incubated for 1 h at 37°C. IP-One-d2 and anti-IP-One Cryptate dissolved in Lysis Buffer (3 μl each) were subsequently added and incubated for 15-30 min at room temperature before measurement. Intracellular IP-One measurement was carried with the IP-One HTRF kit and EnVision multi-plate reader (PerkinElmer) according to the manufacturer’s instructions. Data were normalized to the baseline response of the ligand.

### Molecular docking

Non-standard residues were generated by Discovery Studio 2016 in the Sketch Molecules panel by editing the origin residues correspondingly. Then, the structures encountered a minimization process in Schrödinger Maestro, Protein Preparation Wizard panel. In particular, hydrogens were firstly added to the structure. Then, the protonation state of each residue was assigned with the help of Propka ^63^. Finally, the OPLS3 force field was applied to minimize the energy of the structures with a restrain of heavy atoms to converge them to a root mean square deviation of 0.3 Å.

### Data availability

The atomic coordinates and the electron microscopy maps have been deposited in the Protein Data Bank (PDB) under accession number xxxx, xxxx, xxxx, and xxxx, as well as Electron Microscopy Data Bank (EMDB) accession number xxxx, xxxx, xxxx, and xxxx for the NMU-NMUR1-G_q_-scFv16, NMU-NMUR2-G_q_-scFv16, NMS-NMUR1-G_q_-scFv16, and NMS-NMUR2-G_q_-scFv16 complexes, respectively. Source data are provided with this paper.

### Statistics

All functional study data were analyzed using GraphPad Prism 8.0 (Graphpad Software Inc.) and showed as means ± S.E.M. from at least three independent experiments in triplicate. The significance was determined with two-side, one-way ANOVA with Tukey’s test, and *P* < 0.05 was considered statistically significant.

