## Supplementary files for "Structural insights into the peptide selectivity and activation of human neuromedin U receptors"

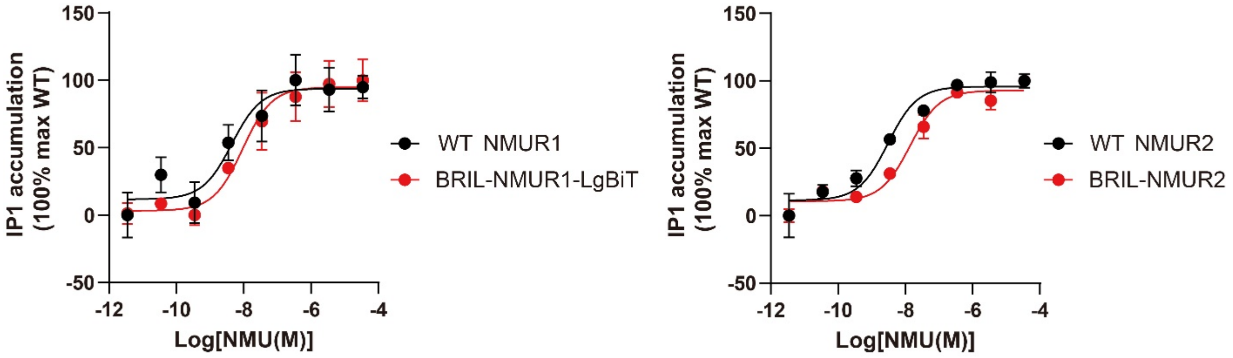


**Supplementary Fig. 1 The comparison of wild-type or engineered NMU receptors response for NMU.** BRIL-NMUR1-LgBiT and BRIL-NMUR2 are constructs used in cryo-EM structure determination. $pEC_{50}$ values of NMU for the wild-type (WT) and modified NMUR1 are 8.35 ± 0.34 and 8.03 ± 0.27, respectively. $pEC_{50}$ values of NMU for the WT and modified NMUR2 are 8.52 ± 0.17 and 7.86 ± 0.14, respectively. Each point represents mean ± S.E.M. from three independent experiments.

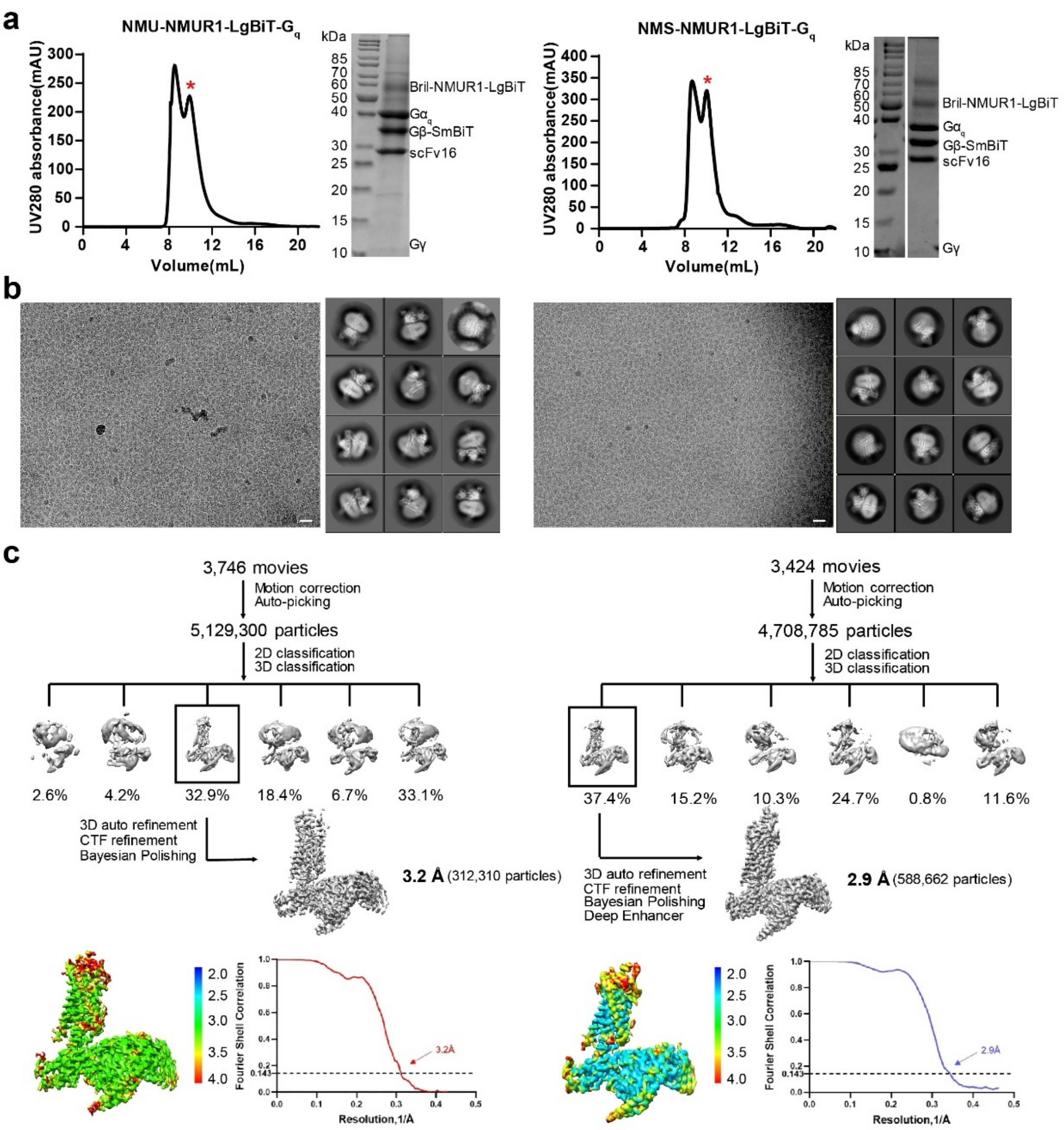


**Supplementary Figure 2. NMU/NMS-NMUR1-$G_q$ complexes purification and cryo-EM data processing. a** Representative size-exclusion chromatography elution profile and SDS-PAGE analysis of NMU-NMUR1-$G_q$ (left) and NMS-NMUR1-$G_q$ complexes (right), respectively. Red asterisks refer to complex monomer. **b** Cryo-EM micrographs and representative 2D average classes of both complexes are shown. Scale bar, 50 nm. **c** Flowchart of cryo-EM data processing, cryo-EM maps (colored by local resolution (Å) calculated using the ResMap package), and "Gold-standard" FSC curves.

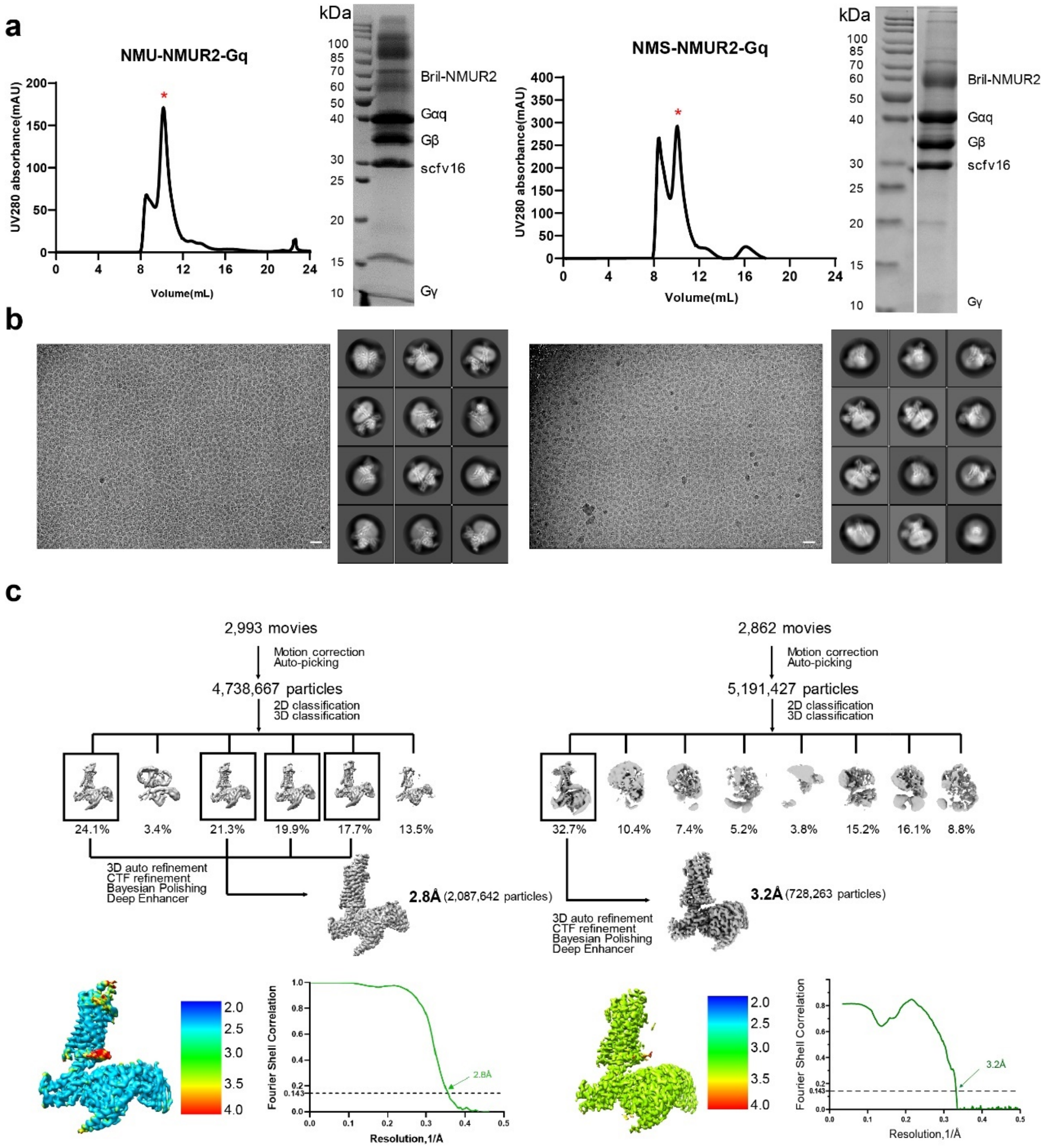


**Supplementary Figure 3. NMU/NMS-NMUR2-$G_q$ complexes purification and cryo-EM data processing. a** Representative size-exclusion chromatography elution profile and SDS-PAGE analysis of NMUR2-NMU-$G_q$ (left) and NMUR2-NMS-$G_q$ (right) complexes, respectively. **b** Cryo-EM micrographs and representative 2D average classes of both complexes are shown. Scale bar, 50 nm. Red asterisks refer to complex monomer. **c** Flowchart of cryo-EM data processing, cryo-EM maps (colored by local resolution (Å) calculated using the Resmap package), and “Gold-standard” FSC curves.

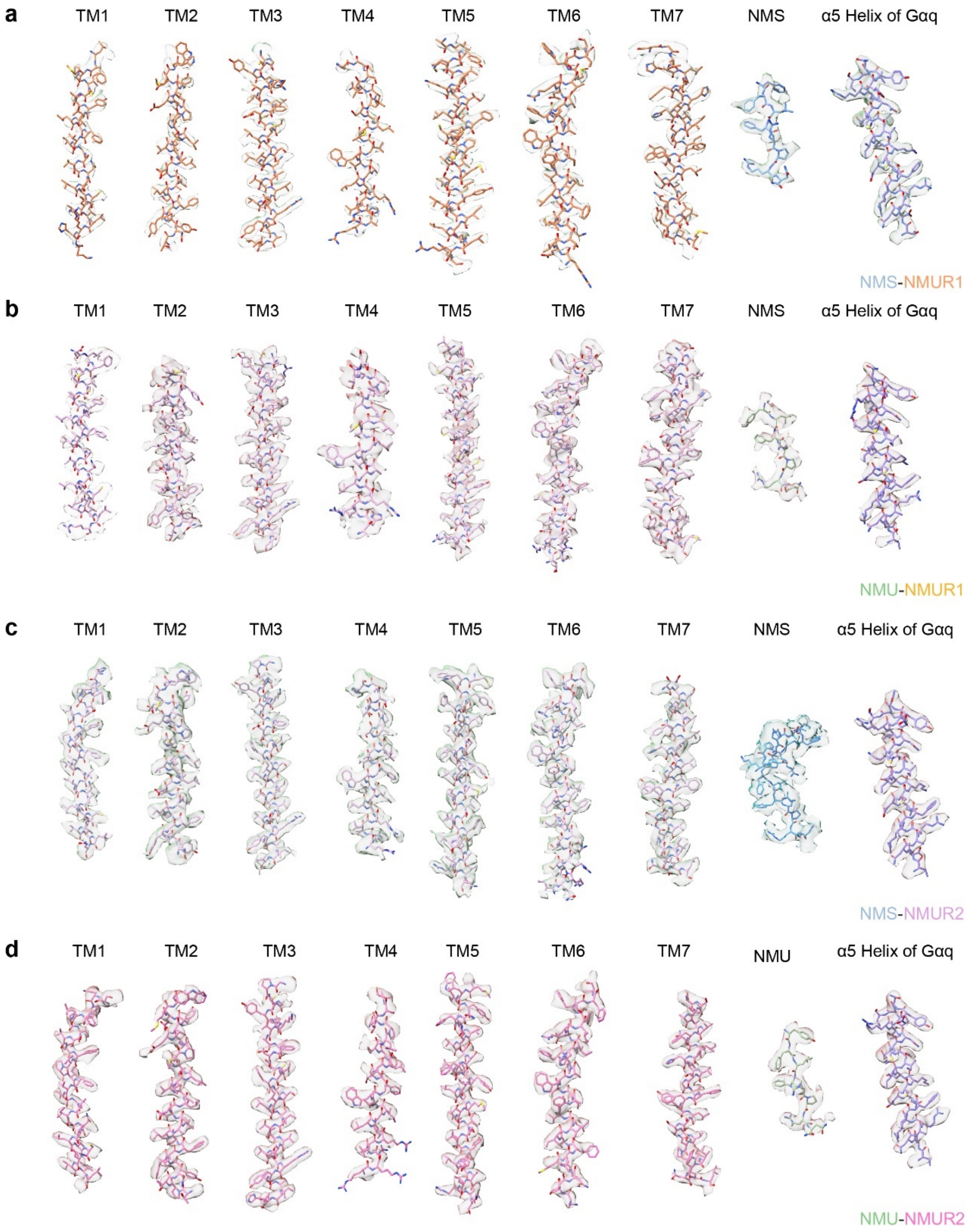


**Supplementary Figure 4. Representative cryo-EM density maps of the NMU/NMS-NMUR1-$G_q$ and NMU/NMS-NMUR2-$G_q$ complexes.** Cryo-EM density maps of the seven transmembrane (TM) helices, α5 helix of $G\alpha_q$, and corresponding peptides for NMS-bound NMUR1 (**a**), NMU-bound NMUR1 (**b**), NMS-bound NMUR2 (**c**), and NMU-bound NMUR2 (**d**) were shown. Colors are shown as indicated.

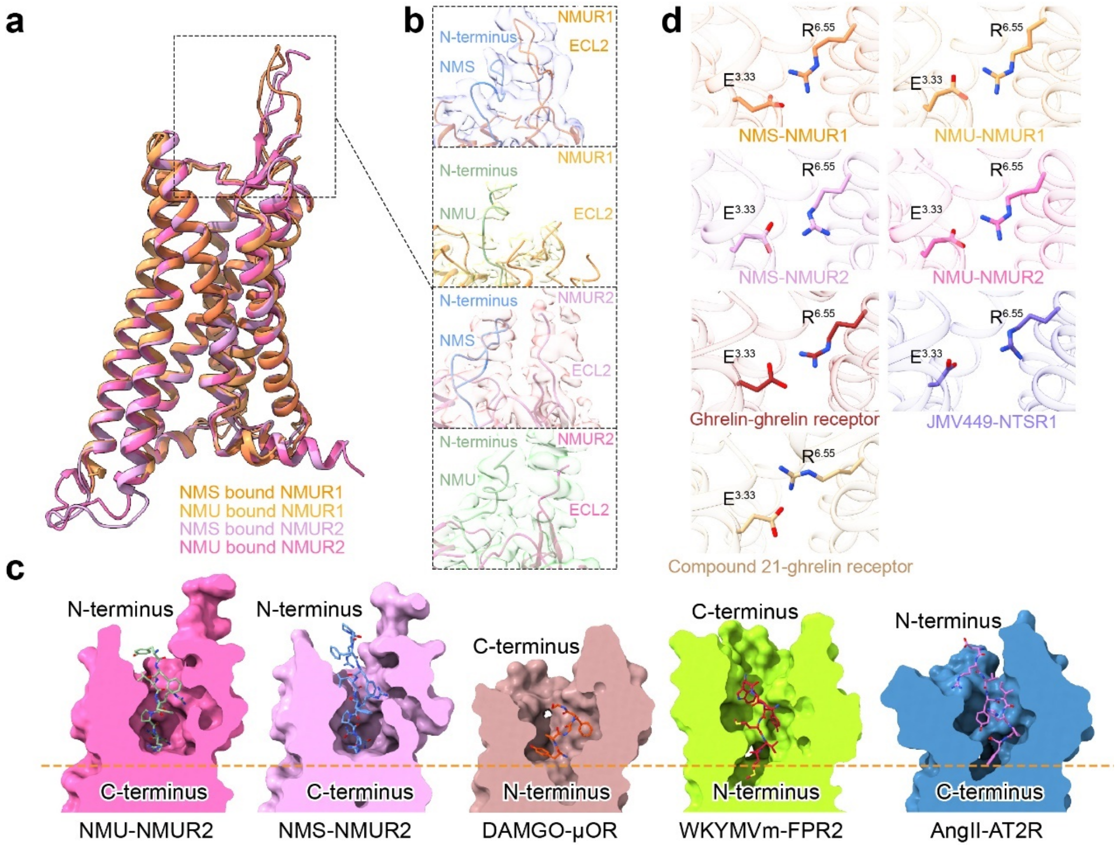


**Supplementary Figure 5. Structural comparison of active NMURs with other peptide-bound class A GPCRs. a** Structural superposition of four NMURs bound to NMU and NMS, respectively. **b** The possible interaction between N-termini of NMU/NMS and ECL2 of receptors. Ambiguous EM densities of the N-termini of peptides can be observed in these four complexes. **c** Structural comparison of the overall peptide-binding pocket of NMUR2 with other GPCRs solved to date. The orange dashed line refers to the bottom of ligand binding pockets of NMUR2. μOR, Mu Opioid Receptor (PDB 6DDE); FPR2, Formyl peptide receptor 2 (PDB 6OMM); AT2R, Angiotensin II type 2 receptor (PDB 6JOD). **d** The conserved salt bridge formed between E/D$^{3.33}$ and R$^{6.55}$. Colors are shown as indicated. Ghrelin receptor (ghrelin-bound, PDB 7F9Y); NTSR1, Neurotensin receptor type 1 (PDB 6OS9); Ghrelin receptor (compound 21-bound, PDB 6KO5).

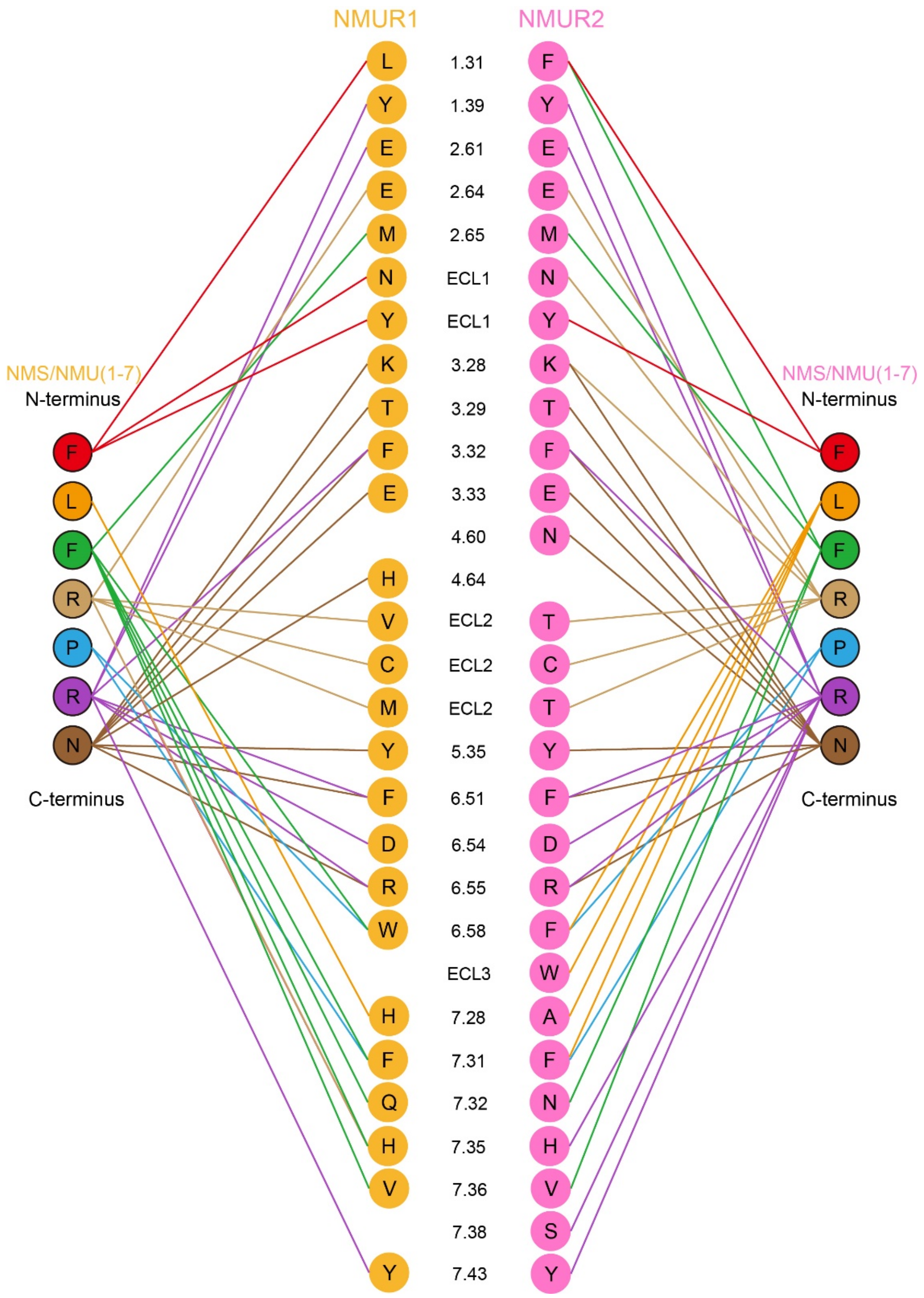


**Supplementary Figure 6. Representative peptide-receptor interaction networks of NMU/NMS-NMUR1-$G_q$ and NMU/NMS-NMUR2-$G_q$ complexes.** Colors are shown as indicated.

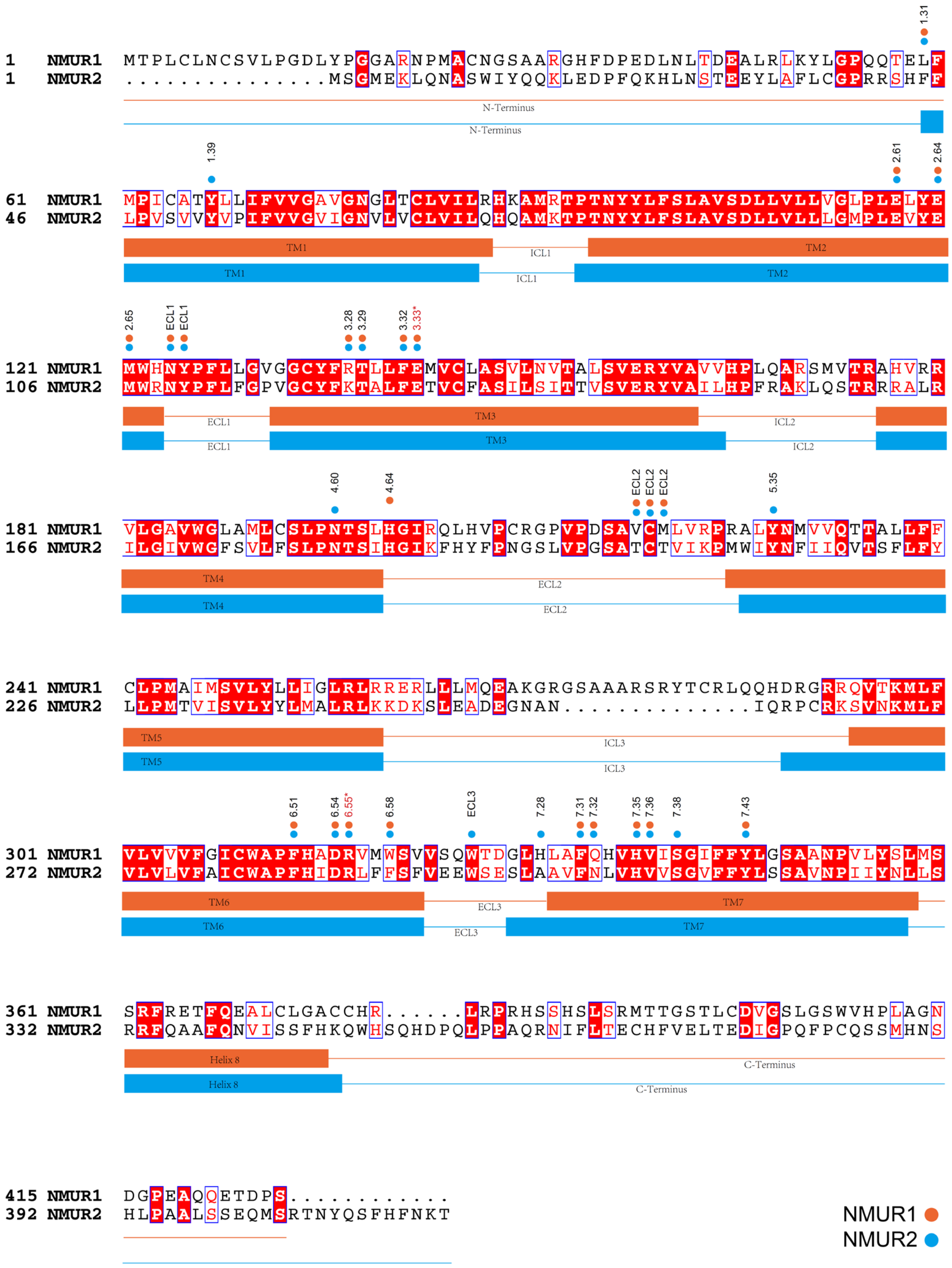

**Supplementary Figure 7. Sequence alignment of the NMUR subfamily.** The sequence alignment of NMUR1 and NMUR2 was generated by CLUSTALW (https://www.genome.jp/tools-bin/clustalw) and ESPript 3.0 (https://espript.ibcp.fr/ESPript/cgi-bin/ESPript.cgi). α-helices for NMUR1 and NMUR2 are shown as columns underneath the sequence. Orange dots represent the binding-pocket residues of NMUR1 bound to NMU/NMS. Blue dots represent the binding pocket residues of NMUR2 bound to NMU/NMS.

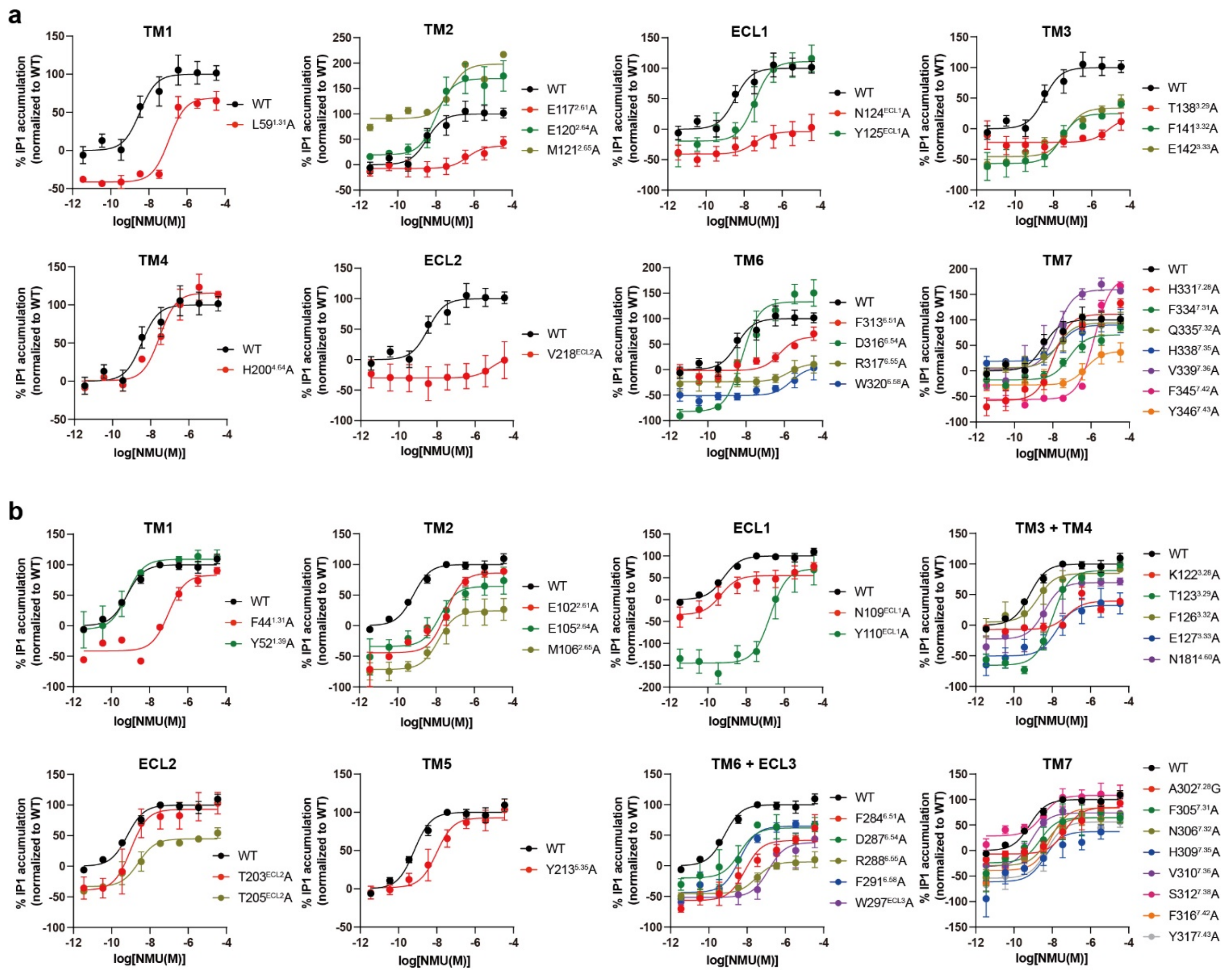


**Supplementary Figure 8. Effects of mutation in the ligand-binding pocket on IP-One accumulation.** AD293 cells were transfected with wild-type (WT), NMUR1, or NMUR2 mutant constructs. Intracellular IP1 accumulation signals were monitored after stimulation with NMU. Each point represents mean ± S.E.M. from three independent experiments. The dataset links to Supplementary Fig. 8 and Supplementary Table 2.

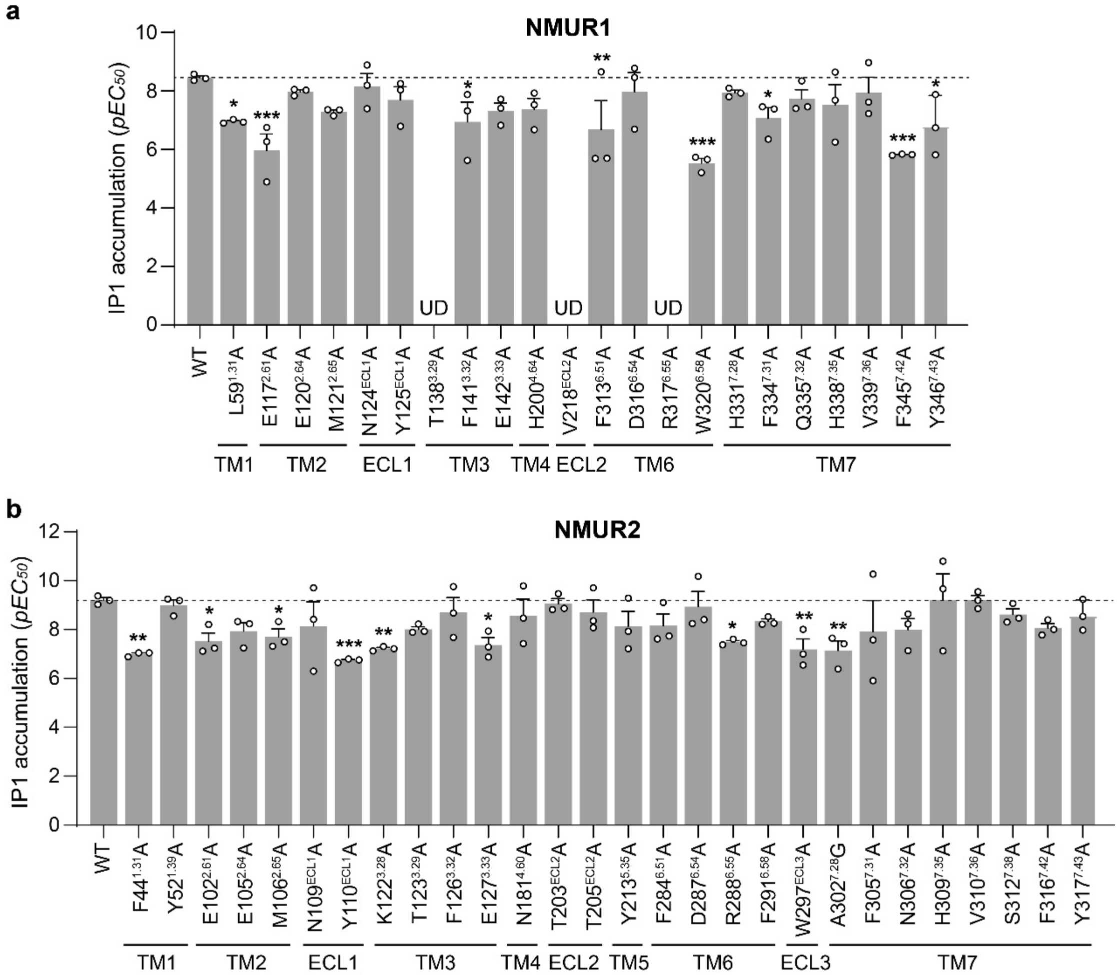


**Supplementary Figure 9. Alanine mutagenesis analysis of peptide-binding pocket of NMURs.** The IP-One assay was performed to evaluate the effects of NMU on the $G_q$-coupling activity of NMUR1 (**a**) and NMUR2 (**b**) mutants. Data were shown as mean $pEC_{50}$ ± S.E.M. from three independent experiments in triplicate (n=3). The significance was determined with two-side, one-way ANOVA with Tukey's test. **P* < 0.05, ***P* < 0.01, ****P*<0.001 *vs.* wild-type (WT) receptor. UD, Undetectable. The dataset links to Supplementary Fig. 7 and Supplementary Table 2.

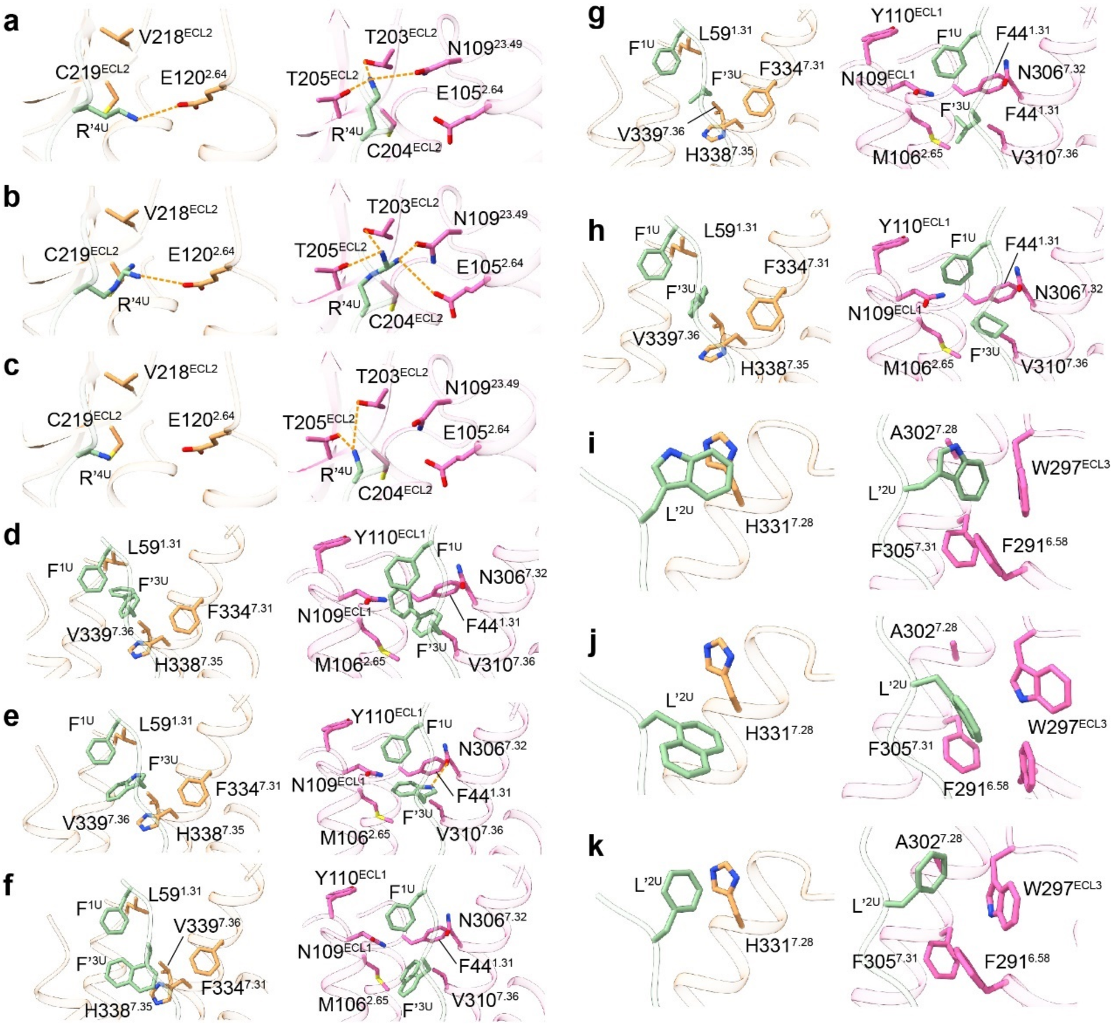


**Supplementary Figure 10.** Molecular docking analysis of selected NMU derivatives for NMUR1 and NMUR2. **a-c** Detailed interaction of various modified $R^4$ of NMU with residues in NMUR1 (Left) and NMUR2 (Right). (**a**) the guanidinium group was replaced by amino, (**b**) the side chain was shortened by one carbon atom, (**c**) guanidine replacement by amino and side chain shortened by two carbon atoms. **d-h** Detailed interaction of various modified $F^4$ of NMU with residues in NMUR1 (Left) and NMUR2 (Right). Biphenyl (**d**), indolyl (**e**), naphthyl (**f**), isopropyl (**g**) and cyclohexyl (**h**) substitutions were introduced, respectively. **i-k** Detailed interaction of various modified $L^2$ of NMU with residues in NMUR1 (Left) and NMUR2 (Right). Indolyl (**i**), naphthyl (**j**), and naphthyl (**k**) replacement were applied, respectively. Non-standard residues were generated by Discovery Studio 2016. The structures encountered a minimization process in Schrödinger Maestro. Colors are shown as indicated. Hydrogen bonds and salt bridges are depicted as orange dashed lines.

a

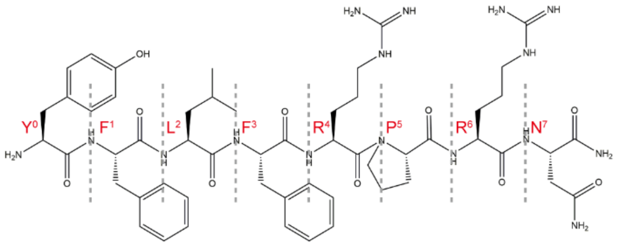


NMU-8 (Lead peptide)
$EC_{50}$(NMUR1)= 0.11 nM
$EC_{50}$(NMUR2)= 0.3 nM

b

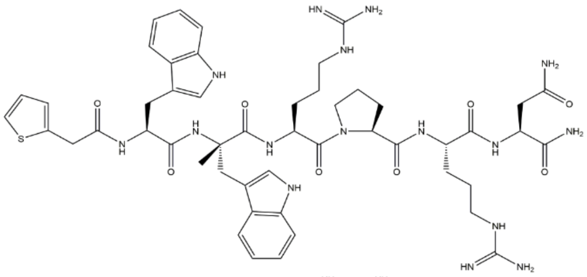

CPN-267
$EC_{50}$(NMUR1)= 0.25 nM
$EC_{50}$(NMUR2)= below $10^{-7}$ M

c

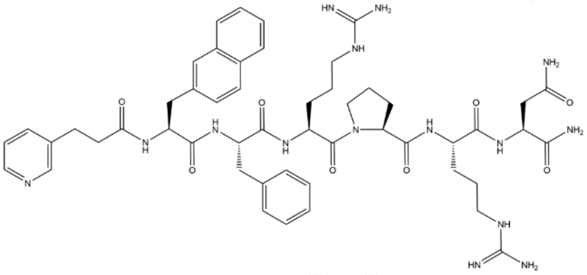

compound 8d (PMID:24999562)
$EC_{50}$(NMUR1)= 5.1 nM
$EC_{50}$(NMUR2)= 249 nM

d

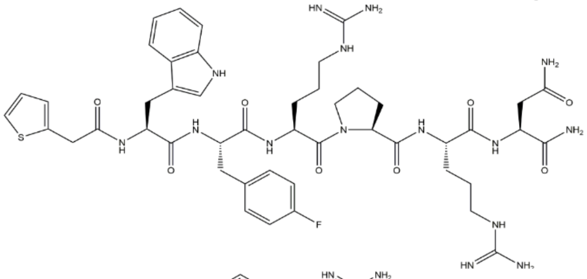

compound 5d (PMID:25815150)
$EC_{50}$(NMUR1)= 0.083 nM
$EC_{50}$(NMUR2)= 2.6 nM

e

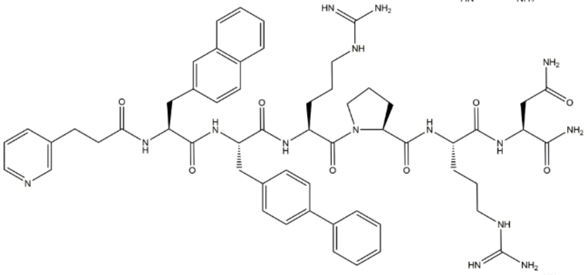

CPN-223
$EC_{50}$(NMUR1)= 3.2 nM
$EC_{50}$(NMUR2)= >1000 nM

f

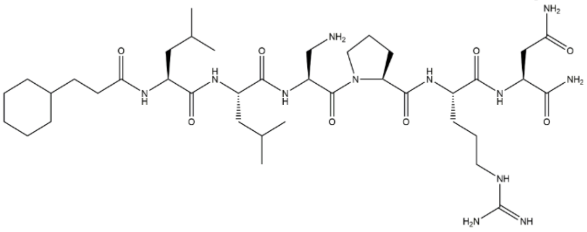

compound 6b (PMID:24999562)
$EC_{50}$(NMUR1)= >1000 nM
$EC_{50}$(NMUR2)= 6.6 nM

**Supplementary Figure 11. Summary of reported NMUR1 and NMUR2 selective agonists derived from NMU-8.**

**Supplementary Table 1. Cryo-EM data collection, model refinement and validation statistics.**

| | NMU-NMUR1-$G_q$ (EMD-32313) (PDB 7W53) | NMS-NMUR1-$G_q$ (EMD-32315) (PDB 7W56) | NMU-NMUR2-$G_q$ (EMD-32314) (PDB 7W55) | NMS-NMUR2-$G_q$ (EMD-32316) (PDB 7W57) |
|---|---|---|---|---|
| **Data collection and processing** | | | | |
| Magnification | 64,000 | 64,000 | 64,000 | 64,000 |
| Voltage (kV) | 300 | 300 | 300 | 300 |
| Electron exposure (e–/Å$^2$) | 61.8 | 61.8 | 61.8 | 61.8 |
| Defocus range (μm) | -1.0~-3.0 | -1.0~-3.0 | -1.0~-3.0 | -1.0~-3.0 |
| Pixel size (Å) | 1.08 | 1.08 | 1.08 | 1.08 |
| Symmetry imposed | C1 | C1 | C1 | C1 |
| Initial particle images (no.) | 5,129,300 | 4,708,785 | 4,738,667 | 5,191,427 |
| Final particle images (no.) | 312,310 | 588,662 | 2,087,642 | 728,263 |
| Map resolution (Å) | 3.2 | 2.9 | 2.8 | 3.2 |
| FSC threshold | 0.143 | 0.143 | 0.143 | 0.143 |
| Map resolution range (Å) | 2-4 | 2-4 | 2-4 | 2-4 |
| **Refinement** | | | | |
| Initial model used (PDB code) | | | | |
| Model resolution (Å) | 3.3 | 3.2 | 3.1 | 3.1 |
| FSC threshold | 0.5 | 0.5 | 0.5 | 0.5 |
| Model resolution range (Å) | 50-3.3 | 50-3.2 | 50-3.1 | 50-3.1 |
| Map sharpening *B* factor (Å$^2$) | -100 | -77.9806 | -199.894 | -172.943 |
| Model composition | | | | |
| Non-hydrogen atoms | 8709 | 8880 | 9220 | 9150 |
| Protein residues | 1109 | 1145 | 1167 | 1167 |
| Ligands | - | - | - | - |
| *B* factors (Å$^2$) | | | | |
| Protein | 51.43 | 61.24 | 54.33 | 45.26 |
| Ligand | - | - | - | - |
| R.m.s. deviations | | | | |
| Bond lengths (Å) | 0.005 | 0.007 | 0.006 | 0.004 |
| Bond angles (°) | 0.650 | 0.665 | 1.021 | 0.581 |
| Validation | | | | |
| MolProbity score | 1.51 | 1.45 | 1.29 | 1.25 |
| Clashscore | 5.02 | 5.76 | 4.80 | 4.25 |
| Poor rotamers (%) | 0.21 | 0.61 | 0.40 | 0.89 |
| Ramachandran plot | | | | |
| Favored (%) | 96.39 | 97.24 | 97.82 | 97.82 |
| Allowed (%) | 3.61 | 2.76 | 2.18 | 2.18 |
| Disallowed (%) | 0.00 | 0.00 | 0.00 | 0.00 |

**Supplementary Table 2.** *$pEC_{50}$* values of NMU on NMUR1 and NMUR2 mutants. IP-One assay was performed to evaluate NMU-induced receptor activation. Data are presented as means ± S.E.M. of three independent experiments (n=3). All data were analyzed by two-side, one-way ANOVA with Tukey's test. *$P$<0.05, **$P$<0.01, ***$P$<0.001 *vs.* wild-type (WT). The dataset links to Supplementary Figs. 7 and 8.

| BW Numbering | NMUR1 Mutant | IP1 accumulation $pEC_{50}$ ± S.E.M. | Surface expression (%WT) ± S.E.M | NMUR2 Mutant | IP1 accumulation $pEC_{50}$ ± S.E.M. | Surface expression (%WT) ± S.E.M |
|---|---|---|---|---|---|---|
| - | WT | 8.45 ± 0.06 | 100 | WT | 9.20 ± 0.10 | 100 |
| 1.31 | L59A | 6.96 ± 0.04 * | 119 ± 11 | F44A | 6.99 ± 0.05 ** | 144 ± 21 |
| 1.39 | NT | NT | NT | Y52A | 9.00 ± 0.22 | 114 ± 17 |
| 2.61 | E117A | 5.97 ± 0.56 *** | 44 ± 3 | E102A | 7.51 ± 0.35 * | 69 ± 4 |
| 2.64 | E120A | 7.97 ± 0.07 | 78 ± 3 | E105A | 7.93 ± 0.35 | 192 ± 4 |
| 2.65 | M121A | 7.29 ± 0.07 | 151 ± 14 | M106A | 7.71 ± 0.31 * | 240 ± 4 |
| ECL1 | N124A | 8.16 ± 0.43 | 80 ± 1 | N109A | 8.14 ± 0.99 | 171 ± 8 |
| ECL1 | Y125A | 7.70 ± 0.45 | 87 ± 3 | Y110A | 6.73 ± 0.05 *** | 45 ± 2 |
| 3.28 | NT | NT | NT | K122A | 7.21 ± 0.05 ** | 42 ± 1 |
| 3.29 | T138A | UD | 110 ± 27 | T123A | 8.01 ± 0.11 | 250 ± 27 |
| 3.32 | F141A | 6.94 ± 0.67 * | 149 ± 12 | F126A | 8.70 ± 0.60 | 113 ± 9 |
| 3.33 | E142A | 7.32 ± 0.26 | 96 ± 11 | E127A | 7.36 ± 0.31 * | 193 ± 19 |
| 4.60 | NT | NT | NT | N181A | 8.56 ± 0.68 | 214 ± 55 |
| 4.64 | H200A | 7.38 ± 0.37 | 95 ± 18 | NT | NT | NT |
| ECL2 | V218A | UD | 142 ± 7 | T203A | 9.05 ± 0.21 | 172 ± 50 |
| ECL2 | NT | NT | NT | T205A | 8.71 ± 0.50 | 67 ± 7 |
| 5.35 | NT | NT | NT | Y213A | 8.14 ± 0.61 | 121 ± 16 |
| 6.51 | F313A | 6.68 ± 0.98 ** | 97 ± 8 | F284A | 8.15 ± 0.48 | 154 ± 13 |
| 6.54 | D316A | 7.98 ± 0.65 | 156 ± 53 | D287A | 8.94 ± 0.62 | 62 ± 5 |
| 6.55 | R317A | UD | 74± 4 | R288A | 7.48 ± 0.07 * | 130 ± 1 |
| 6.58 | W320A | 5.54 ± 0.16 *** | 81 ± 11 | F291A | 8.33 ± 0.10 | 189 ± 14 |
| ECL3 | NT | NT | NT | W297A | 7.18 ± 0.43 ** | 76 ± 4 |
| 7.28 | H331A | 7.94 ± 0.08 | 111 ± 30 | A302G | 7.15 ± 0.39 ** | 172 ± 50 |
| 7.31 | F334A | 7.07 ± 0.36 * | 113 ± 1 | F305A | 7.91 ± 1.27 | 87 ± 3 |
| 7.32 | Q335A | 7.74 ± 0.31 | 124 ± 12 | N306A | 8.00 ± 0.45 | 167 ± 10 |
| 7.35 | H338A | 7.53 ± 0.69 | 222 ± 22 | H309A | 9.19 ± 1.08 | 174 ± 25 |
| 7.36 | V339A | 7.94 ± 0.53 | 97 ± 3 | V310A | 9.20 ± 0.20 | 83 ± 6 |
| 7.38 | NT | NT | NT | S312A | 8.61 ± 0.23 | 140 ± 8 |
| 7.42 | F345A | 5.83 ± 0.01 *** | 121 ± 4 | F316A | 8.06 ± 0.18 | 112 ± 7 |
| 7.43 | Y346A | 6.81 ± 0.60 * | 98 ± 4 | Y317A | 8.58 ± 0.36 | 126 ± 6 |

BW numbering, Ballesteros & Weinstein numbering, a generic GPCR residue numbering scheme; UD, Undetectable; NT, Not tested.
